# Rewiring of mite allergen-specific Th-memory-associated gene networks during immunotherapy

**DOI:** 10.1101/473561

**Authors:** Anya C. Jones, Denise Anderson, Niamh M. Troy, Dominic Mallon, Rochelle Hartmann, Michael Serralha, Barbara Holt, Anthony Bosco, Patrick G. Holt

## Abstract

**Background:** Multiple regulatory mechanisms have been identified employing conventional hypothesis-driven approaches as contributing to allergen-specific immunotherapy outcomes, but understanding of how these integrate to maintain immunological homeostasis is incomplete.

**Objective:** To explore the potential for unbiased systems-level gene co-expression network analysis to advance understanding of immunotherapy mechanisms.

**Methods:** We profiled genome-wide allergen-specific Th-memory-associated responses prospectively during 24mths subcutaneous immunotherapy (SCIT) in 25 rhinitics, documenting changes in immunoinflammatory pathways and associated co-expression networks and their relationships to symptom scores out to 36mths.

**Results:** Prior to immunotherapy, mite-specific Th-memory-associated response networks involved multiple discrete co-expression modules including those related to Th2-, Type1-IFN-, Inflammation-, and FOXP3/IL2-associated signalling. A signature comprising 109 genes correlated with symptom scores, and these mapped to cytokine signalling/T-cell activation-associated pathways, with upstream drivers including hallmark Th1/Th2-and inflammation-associated genes. Reanalysis after 3.5mths SCIT updosing detected minimal changes to pathway/upstream regulator profiles despite 32.5% symptom reduction, however network analysis revealed underlying merging of FOXP3/IL2-with Inflammation-and Th2-associated modules. By 12mths SCIT, symptoms had reduced by 41% without further significant changes to pathway/upstream regulator or network profiles. Continuing SCIT to 24mths stabilised symptoms at 47% of baseline, accompanied by upregulation of the Type1-IFN-associated network module and its merging into the Th2/FOXP3/IL2/Inflammation module.

**Conclusions:** SCIT stimulates progressive integration of Th-memory-associated Th2-, FOXP3/IL2-, Inflammation-, and finally Type1-IFN-signalling subnetworks, forming a single highly integrated co-expression network module, maximising potential for stable homeostatic control of allergen-specific Th2 responses via cross-regulation. Th2-antagonistic Type1-IFN signalling may play a key role in stabilising clinical effects of SCIT.

## INTRODUCTION

The primary aim of allergen-specific immunotherapy (SIT) is attenuation of downstream inflammation resulting from allergen-specific CD4^+^Th-memory cell activation. Multiple regulatory mechanisms are believed to contribute to SIT-induced modulation of Th-memory-dependent inflammatory symptoms^1-4^, but the understanding of how these operate individually/collectively is relatively scant. Traditional approaches to this question have emphasised hypothesis-driven studies on candidate targets selected based on *a priori* knowledge, focusing on relatively limited numbers of effector/regulatory mechanisms. However, it is now recognised that immune responses, including those induced by allergens^5,6^, involve thousands of genes functioning within complex networks, and hence the application of systems-level analytical methodology which can capture and deconvolute some of this complexity may prove useful in elucidating SIT mechanisms.

In this regard, our group has previously employed genome wide expression profiling of allergen-activated CD4^+^T-cells to define a highly inter-correlated Th2-associated gene subnetwork that is operative in atopic children at risk of allergic disease^5-7^. In the analyses below we have extended this systems-level approach, to identify the full spectrum of functionally coherent gene subnetworks operating within aeroallergen-specific Th-memory-associated responses in highly symptomatic adults, and to map changes in their architecture and interconnectivity during two years of SIT, in particular changes associated with reduction in Th2-associated allergy symptoms.

## METHODS

### Study population

House dust mite (HDM) sensitised subjects (SPT^+^ and/or specific IgE ≥0.35kU/L) with perennial rhinitis (n=25; **Table E1**) were recruited at Fremantle Hospital, Western Australia, prior to commencement of a standard 3yr course of HDM-specific subcutaneous immunotherapy (SCIT). The SCIT protocol and mechanistic study design was approved by the institutional ethics committee, and consent obtained from all subjects.

### Immunotherapy protocol

The protocol was based on Blumberga^8^, utilising Alutard^®^ SQ *Dermataphagoidies pteronyssinus* (ALK, Horsholm, Denmark). Treatment included 15-week updosing (up to 100,000 SQ-U) and 3-year maintenance with injection intervals of 6±2 weeks; patients recorded daily respiratory symptoms and medication use, the details of which were utilised to generate symptom scores at the end of designated treatment periods^9^.

### PBMC processing

PBMC were collected prior to immunotherapy (Visit 1/V1), and at 3.5mths (V2), 12mths (V4) and 24mths into treatment (V5; **Figure E1**), and cryopreserved as described^10^.

### Analytical strategy

This exploratory study aimed to evaluate the potential of network analytic methodology to elucidate short-/long-term effects of SCIT on allergen-specific Th-memory-associated responses. Our approach (**Figure E2**) involved initial *in vitro* culture of unfractionated PBMC with unfractionated HDM allergen extract to enable interactions to occur between allergen-specific and non-specific T-cells and bystander myeloid cells. We have previously demonstrated that the overall HDM-induced Th-memory-associated response signature in atopics comprises components from both HDM-triggered and (cytokine-triggered) HDM-unresponsive CD4+T-cells^5,7^, hence we purified CD4^+^T-cells from these cultures for transcriptomic profiling without additional fractionation. Microarray analysis was used to compute T-cell differentially expressed gene (DEG) signatures, followed by pathway analysis to identify functionally coherent pathways that map to these signatures (IL2-signalling; Th2 etc). This was followed by upstream regulator analysis (URA) to identify “driver” genes in specific Th-memory cells and/or in bystander myeloid/T-cell populations in PBMC, that are statistically likely to have contributed to the 24hr T-memory-associated signatures at an earlier stage in the activation process.

Independently, network analysis was performed to identify gene coexpression networks that underpin allergen-driven Th-memory-associated responses. A subset of the subjects also received grass allergens in the SCIT mix, and sub-analysis of study outcome samples after stratification by treatment to ascertain the effects of the latter on HDM-induced DEG profiles.

### *In vitro* cell cultures and CD4+T cell isolation

PBMC from visits V1-V4 were thawed and cultured together employing the same batch of LPS-depleted whole HDM extract (10ug/ml; gift from ALK Abello) or medium control for 24h^10^, whilst V5 was cultured as a stand-alone experiment employing the same reagent batches. We have previously established that the AIM-V serum-free medium employed does not support PBMC activation by LPS, including when spiking cultures with up to 100ug/L LPS^10^. CD4^+^T cells were harvested from cultures using Dynabeads(ThermoFisher) for total RNA extraction with TRIzol(Ambion, Life Technologies) followed by RNAeasy MinElute(Qiagen, Hilde, Germany), and subsequent microarray profiling(Affymetrix PrimeView).

### Antibody measurement

HDM-specific IgE and IgG4 levels were quantified by ImmunoCAP(ThermoFisher)^10^.

### Microarray data pre-processing and differential expression analysis

Raw microarray data are available from the Gene Expression Omnibus repository(GEO Accession #GSE122290). Gene expression data were analysed using R statistical computing software(version 3.4.4). A custom chip description file with updated genome information (primeviewhsentrezgcdf, version 22) was utilised to annotate probe sets to genes^11^. Quality control(QC) of raw microarray data was performed with *ArrayQualityMetri*cs^12^; one case/control subject at V5 was removed based on QC plots. The robust multi-array average algorithm(*RMA*^13^), was used for background correction and quantile normalisation. Non-informative probe sets were filtered out using the proportion of variation accounted for by the first principal component algorithm(*PVAC*>0.4^14^), resulting in 3264 probe sets(V1, V2 and V4) and 3825 probe sets(V5) for downstream statistical analysis.

DEG were identified using *limma*^15^, with a false discovery rate control for multiple testing. Quality weights were included in the linear model with the *arrayWeights*()^15^ function. Paired samples were taken into account with the *duplicateCorrelation()*^15^ function and the model estimates were adjusted for batch effects, which had been identified with PCA plots (**Figure E3**). We tested whether response to stimulation is associated with respiratory symptoms at each visit using *limma* by including both covariates and their interaction in the linear model.

### Network analysis

Prior to network construction, raw expression data were pre-processed separately for each visit with *RMA*^13^ and batch variation was removed with *ComBat*^16^. Batch effect removal was verified by PCA plots (**Figures E4/5**) and differences between HDM-stimulated and control cultures were confirmed and visualised with heatmaps (**Figure E6**). The *PVAC*-filtered data resulted in 4027, 3042, 3912 and 3831 probe sets at V1/V2/V4/V5 respectively, and these probe sets were combined resulting in 5265 genes for network analysis. Co-expression networks were constructed employing *WGCNA(*weighted gene coexpression network analysis)^17^, based on the concept that the organisation and function of a biological system can be represented as a network graph of coexpressed genes. We employed the following parameters for *WCGNA*: signed networks, softPower = 11, Pearson correlation, minimum module size = 50, deepsplit = 0, merge cut height = 0.1. Principal component analysis was performed at each visit to determine the relatedness of the modules. The modules at each time point (V1, V2, V4 and V5) were examined for enrichment with differentially expressed genes (HDM stimulated versus control) obtained with *limma*. Then, HDM-response modules were identified based on median adjusted *p*-values < 0.05 (from the *limma* analysis). Network wiring diagrams were constructed utilising the top-weighted 800 network edges (pairwise gene-gene correlations), which were extracted from the adjacency matrix from the expression data. The networks were graphically depicted using Cytoscape^18^ (version 3.6.1), employing the NetworkAnalyzer tool, parameters: undirected edges, organic layout, map node size to degree: low node values to small size.

### Pathways analysis & Gene ontology analysis

Pathway enrichment and gene ontology analysis were performed with InnateDB version 5.4^19^ based on a hypergeometric distribution and Benjamini & Hochberg corrected *P-values*≤0.05. Public databases utilised included INOH, KEGG, NETPATH, PID, BIOCARTA, REACTOME for pathways analysis and gene ontologies including molecular function, cellular component and biological process.

### Upstream regulator analysis

Putative regulators of the gene expression patterns were identified employing Ingenuity Systems Upstream Regulator Analysis(URA)^20^. Molecular drivers with Benjamini & Hochberg adjusted *P-values*≤0.05 and absolute *Z-scores*≥2.0 were deemed significant. The overlap *P-value* measures enrichment and the *Z-score* measures activation/inhibition of the regulator.

## RESULTS

### Evolution of symptoms over time during SCIT

**Figure 1** illustrates symptom scores(mean±95% CI) at each sampling. The largest drop in symptom scores had occurred by the end of the updosing phase(32.5% reduction; V2/3.5mths) at which point maintenance therapy commenced. The mean symptom score decline reached 37% by V4(12mths) and 51% by V5(24mths), remaining at this level to the end of treatment at V6(36mths). Consistent with the literature^1,2,4^, reductions in symptoms across the first year of SCIT were accompanied by increases in serum HDM-specific IgE and IgG4 (**Figure E7A/B**).

**Figure 1.**
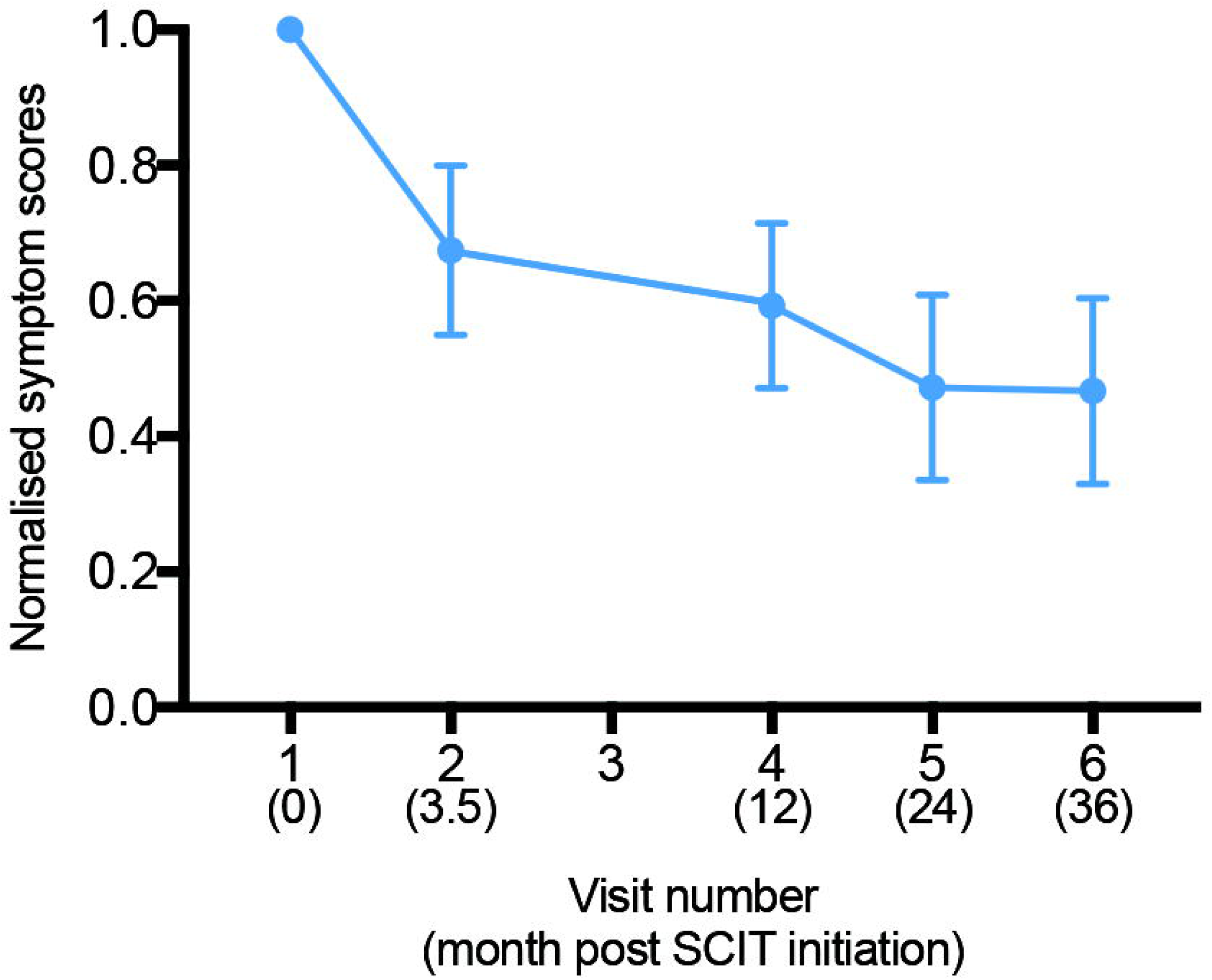
Evolution of symptoms over time during SCIT. Normalised respiratory symptom scores based on symptoms recorded for one week prior to visit dates were plotted (as mean ± 95%CI) for pre-(V1) and post-SCIT treatment (V2/3.5mths, V4/12mths, V5/24mths and V6/36mths sampling points.

### CD4+T cell immune responses pre-treatment

Baseline CD4^+^T-cell responses to HDM at V1 comprised 1715 DEG (**Figure 2A/Tables E2/E3**). Pathways analysis demonstrated enrichment for genes associated with a number of functionally coherent pathways, in particular involving cytokine/chemokine receptor interactions, antigen processing/presentation and T-cell transcriptional activation (**Figure 2B**). URA revealed that the top-ranking putative drivers (ranked in **Figure 2C** by activation *Z-scores*/adjusted *P-values*) were dominated by those associated with Th1/Th2 immunity (including respectively IFNG/IL12; IL4/IL5), inflammation (TNF, IL1B, CSF2, STAT3) and T-cell activation (CD40 ligand, CD3/TCR, CD28).

**Figure 2.**
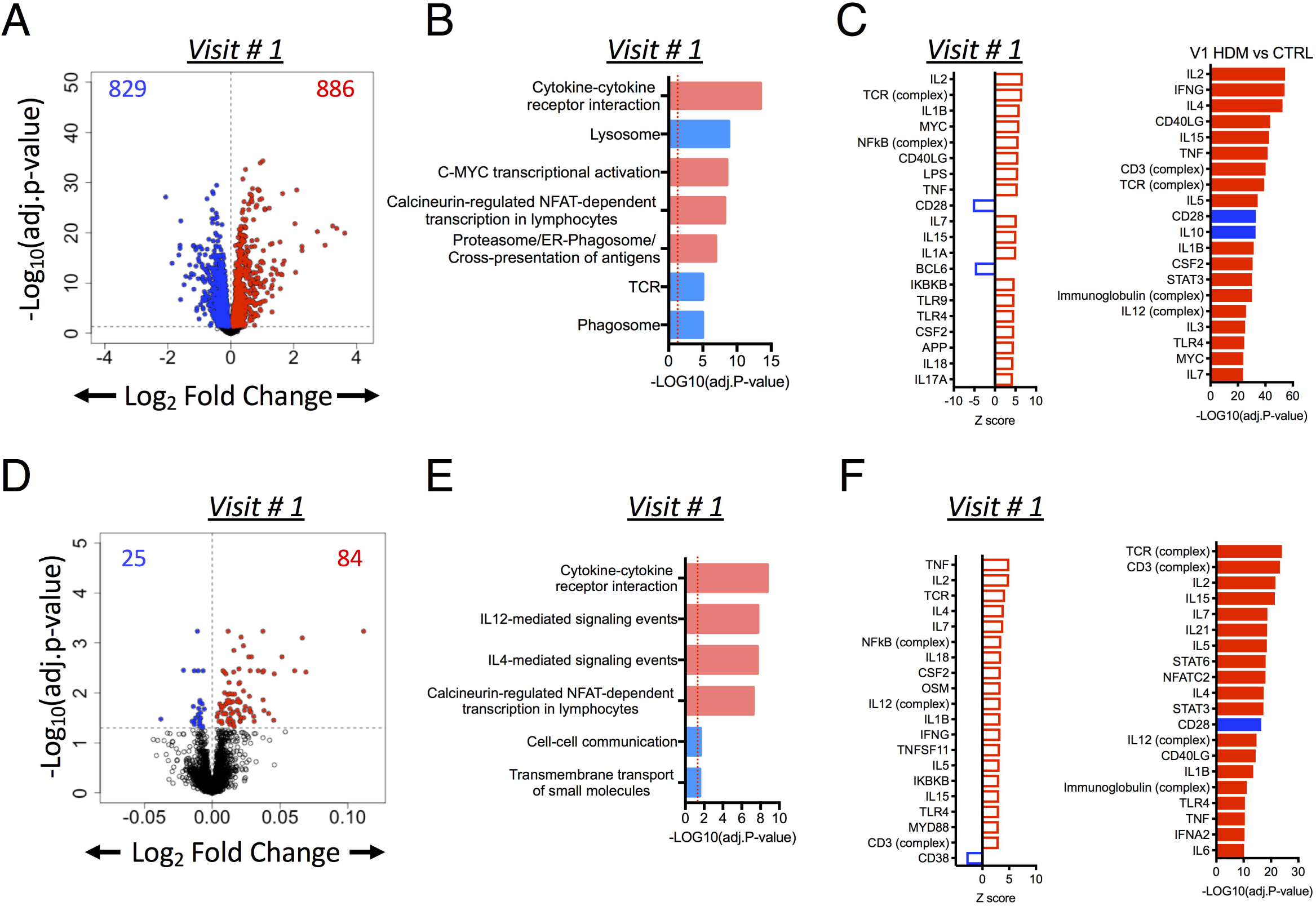
CD4+ T-cell immune responses pre-treatment (V1) were correlated with respiratory symptoms. A) Volcano plot showing HDM-specific DEG at V1 (dashed horizontal line=adjusted *p-value*<0.05; n=25). B) Pathways analysis of DEG was carried out with InnateDB (black=upregulated pathways, open=downregulated pathways). C) Predicted upstream regulators of the DEG (black=activated drivers, open=inhibited drivers). LPS=lipopolysaccharide. D) It was tested if response to stimulation is associated with respiratory symptoms at V1 employing *limma* and volcano plot shows the correlation of respiratory symptoms with CD4+T cell responses at V1 (n=25); E) corresponding up-and-downregulated pathways and F) upstream regulators.

### Association of respiratory symptoms with gene expression pre-treatment

We tested whether allergen-driven T-cell response patterns were associated with respiratory symptom scores at each visit employing *limma*. At pre-treatment(V1) we identified a gene signature comprising 109 DEG as significantly associated with symptoms (**Figure 2D, Table E4**). Pathways analysis showed that symptom-associated genes were enriched for cytokine/cytokine-receptor interactions, Th1-/Th2-mediated signalling, calcineurin-regulated NFAT-dependent lymphocyte transcription, whilst downregulated genes were enriched for cell-cell communication/transmembrane transport (**Figure 2E**). URA demonstrated that the top-ranking drivers were associated with T-cell activation (TCR and CD3 complexes, NFATC2, CD40LG), Th1 immunity (IL12 complex/IFNG), Th2 immunity (IL4/IL5), common gamma chain signalling (IL2, IL4, IL7, IL15, IL21), and inflammation (TNF, STAT3, STAT6, IL1B, IL18; **Figure 2F**).

We then replicated the analysis from **Figure 2D**, in this case seeking to identify gene expression patterns associated cross-sectionally with attenuated post-treatment symptom expression scores at V2, V4 and V5 respectively. No significant associations were detected at these later time points, indicating that the contribution of HDM-specific immunoinflammatory responses to the overall expression of clinical symptoms was significantly diminished from V2 onwards relative to pre-treatment (**Figure E8**).

### Impact of SCIT on HDM-specific Th-memory-associated response profiles

We assessed HDM-induced T-cell response patterns at V2, V4 and V5. In excess of 1800 DEG were detected at each of these sampling times **(Figure 3A, Tables E3, 5-7**). The list of up/downregulated pathways and their putative drivers at the different visits during SCIT (**Figure 3B,C**) closely resembled that identified for V1 (**Figure 2B,C**); the notable exception was the presence of an operational/upregulated type1/2 interferon signalling pathway at V5 (**Figure 3B/C**) that was not active earlier in the time course. In particular, no type1 interferon-associated genes appeared at any level of significance amongst the list of upstream drivers prior to V5. As illustrated in **Table I**, which shows activation *Z-scores* at each visit for the V5 upstream drivers, scores for the interferon-associated drivers are uniformly elevated at V5, in contrast to the stable pattern for other drivers including those related to Th2 inflammation.

**Table I.** Activation Z-scores at each visit for the V5 upstream drivers.

| Upstream regulators | V1 | V2 | V4 | V5 |
| --- | --- | --- | --- | --- |
| NFkB (complex) | 5.80 | 5.85 | 5.84 | 7.05 |
| IL1B | 6.08 | 5.80 | 5.87 | 6.87 |
| CD40LG | 5.74 | 5.94 | 6.04 | 6.50 |
| TCR (complex) | 6.66 | 5.85 | 6.61 | 6.28 |
| IL2 | 6.79 | 6.66 | 6.69 | 6.20 |
| IL7 | 5.32 | 5.78 | 5.27 | 6.14 |
| TNF | 5.58 | 5.12 | 5.32 | 6.10 |
| MYC | 5.97 | 5.75 | 6.36 | 6.07 |
| **IFNA2** | 1.65 | 1.79 | 0.55 | **5.99** |
| **IRF7** | 0.60 | 1.95 | -0.16 | **5.85** |
| **IFNG** | 3.48 | 3.93 | 2.60 | **5.78** |
| TLR4 | 4.71 | 4.80 | 4.11 | 5.62 |
| IL15 | 5.24 | 5.29 | 5.62 | 5.54 |
| MYCN | 4.02 | 4.41 | 4.65 | 5.35 |
| TLR9 | 4.76 | 4.69 | 3.81 | 5.00 |
| **STAT1** | 2.27 | 3.04 | 1.86 | **4.87** |
| IL18 | 4.53 | 4.56 | 4.46 | 4.78 |
| CSF2 | 4.71 | 4.12 | 4.32 | 4.25 |
| IL5 | 4.22 | 4.15 | 4.17 | 4.02 |
| Immunoglobulin (complex) | 2.85 | 3.37 | 3.71 | 3.96 |
| CD3 (complex) | 3.52 | 3.62 | 3.68 | 3.96 |
| IL3 | 3.67 | 3.63 | 3.39 | 3.86 |
| STAT3 | 3.32 | 3.37 | 3.04 | 2.65 |
| IL27 | 1.96 | 1.79 | 1.00 | 2.28 |
| IL4 | 2.49 | 2.44 | 2.53 | 2.19 |
| IL10 | -2.23 | -2.48 | -2.81 | -2.85 |
| CD28 | -5.38 | -4.97 | -5.14 | -4.49 |
| BCL6 | -4.91 | -4.93 | -4.82 | -4.94 |
| RICTOR | -3.64 | -4.95 | -4.35 | -5.14 |
| OSM | 3.14 | 2.85 | 3.04 | 4.63 |

Putative regulators of the DEG were identified at each visit employing Ingenuity Systems Upstream Regulator Analysis. Molecular drivers with Benjamini & Hochberg adjusted *P-values*≤0.05 and absolute *Z-scores*≥2.0 were deemed significant (positive *Z-scores*=activation, negative *Z-scores*=inhibition).

**Figure 3.**
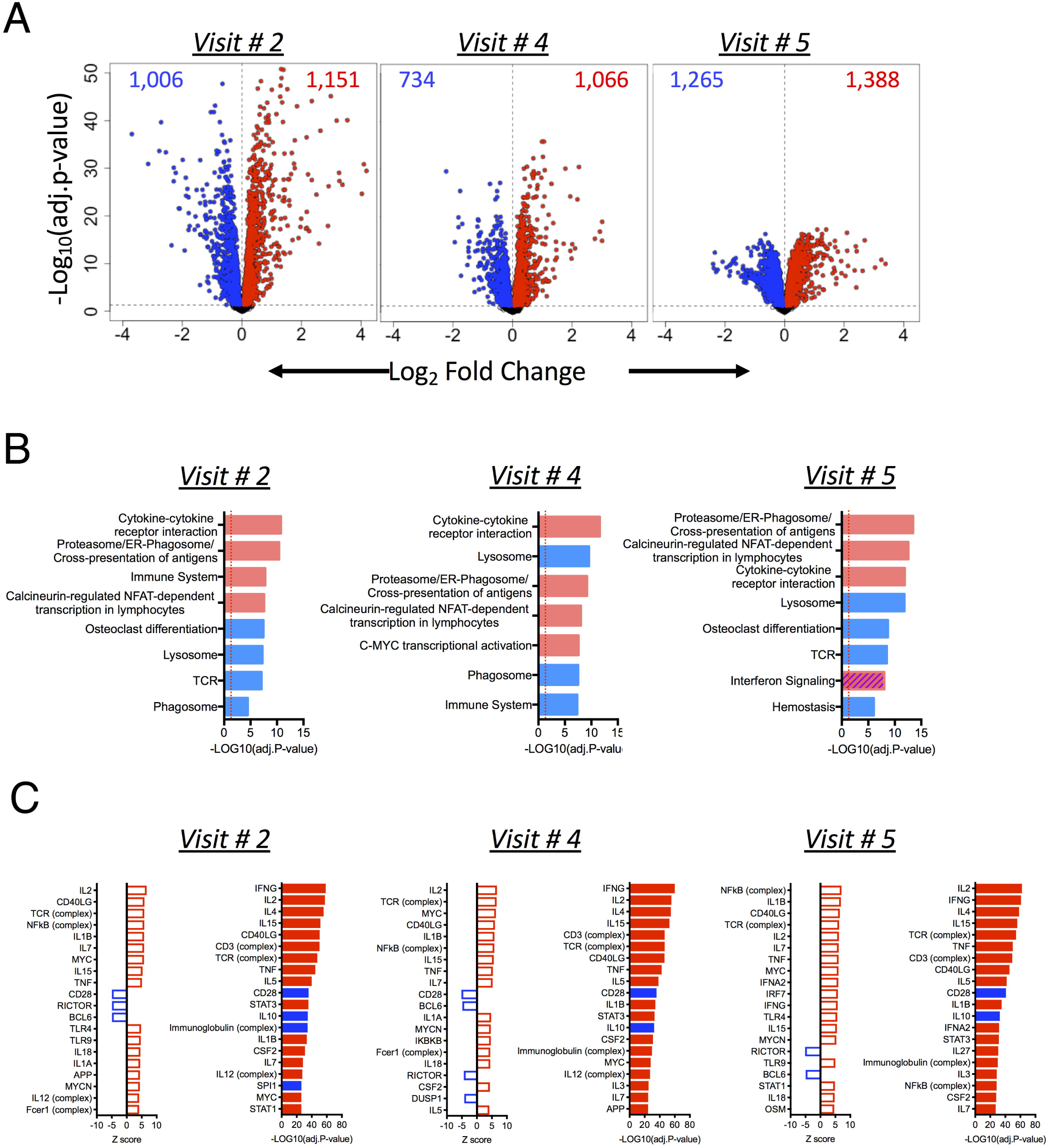
CD4+ T-cell responses over the course of SCIT treatment (V2, V4 and V5). A) Volcano plots showing HDM-specific CD4+ T-cell responses at V2, V4 and V5 respectively (n=25). The dashed horizontal line indicates an adjusted *p-value*<0.05. B) Pathways analysis of DEG was carried out with InnateDB (black=upregulated pathways[includes IFN signaling/striped at V5], open=downregulated pathways). C) Predicted upstream regulators of the DEG (black=activated drivers, open= inhibited drivers).

### *WGCNA* analysis of the HDM-specific T-cell responses

We next constructed weighted gene co-expression networks at each sampling to obtain a systems-level understanding of how the correlation patterns between genes changed over the course of allergen-specific immunotherapy. First, we calculated ranked expression levels and ranked network connectivity, and compared the data at V1 with subsequent visits. As illustrated in **Figure E9**, the data showed that variations in network connectivity (i.e. gene-gene co-expression patterns) were most striking at V2 and V5. These findings were reflected by conventional statistical analyses, which identified hundreds of DEG between the respective responses at V2 versus V1, and V5 versus V1 (Tables E7/E8). Variations in network connectivity were also evident at V4, even though no differentially expressed genes were identified between the respective responses at V1 and V4.

We next looked for changes in modular architecture of the co-expression networks during SCIT. To illustrate this, we superimposed the network connectivity patterns for each visit over the modular architecture from V1 (**Figure 4A**). The data showed that V2 and V5 were characterised by the presence of strong correlation patterns off the diagonal, suggesting rewiring between modules. Moreover, this was also reflected by changes in the modular architecture from V1 to V5 (indicated by colours below **Figure 4B**, left panel; and the roadmap showing the progressive re-wiring of modules (denoted A-N) at ensuing visits (**Figure 4B**, right panel). The co-expression networks were organised into 9/7/10/6 modules at V1/V2/V4/V5 respectively (**Figure 4B**). We then interrogated the modules for functional coherence and examined them for DEG enrichment (**Figure 4C**). At pre-immunotherapy V1, the modules denoted C/F/G/I, relating respectively to metabolism of proteins/Th2 immunity/inflammation/IL2-signalling, were upregulated. Of note, type1 interferon-associated genes formed a discrete module(A), however, these were not differentially expressed in the response. Following completion of updosing at V2, the modular architecture had become restructured and the modules F, G and I amalgamated forming one upregulated module J (**Figure 4B,C**), whilst the type1 interferon network remained unchanged, and lysosome/phagosome/antigen-processing-presentation module C and TCR signalling module D were downregulated.

**Figure 4.**
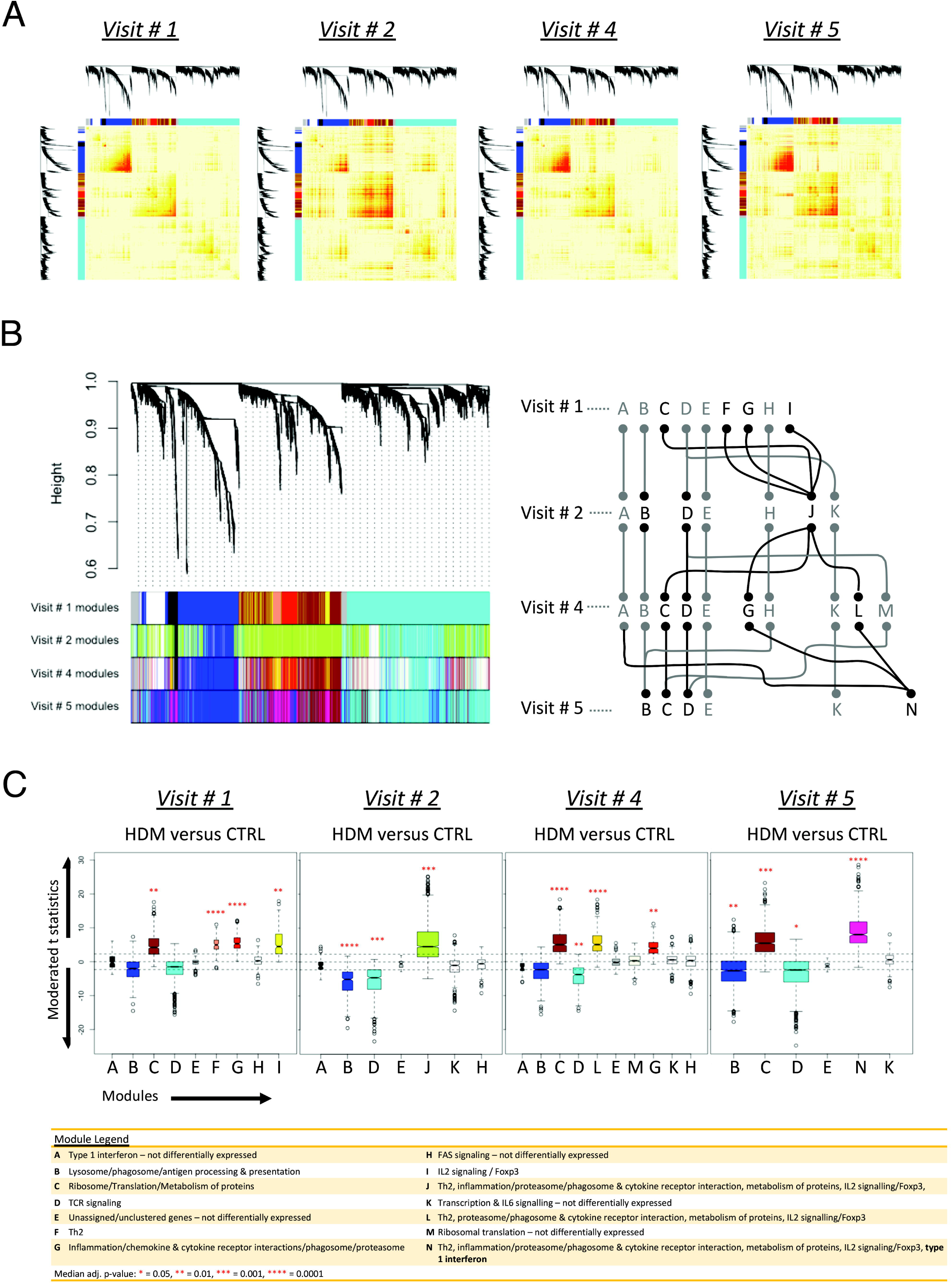
Network analysis of the HDM-induced T cell responses. Co-expression networks were constructed employing *WGCNA* (n=25, V1/V2/V4; n=24, V5). A) Network connectivity patterns(V2/V4/V5) were superimposed over V1 modular architecture. B) Comparison of V1 modular architecture with the module assignments V2/V4/V5 (left panel). Roadmap showing progressive re-wiring of the modules (A-N) over the course of SCIT(right panel; black circles/lines=differentially expressed, grey=not differentially expressed. C) Box-and-whisker plots showing modules with DEG. Median adjusted *p-values:* * <0.05,** <0.01,*** <0.001,**** <0.0001.

At V4, the IL2 signalling and Th2 networks remained fused (module L) while the inflammation/lysosome/phagosome/antigen processing-presentation (G), metabolism of proteins (C), and type1 interferon-associated (A) modules remained as discrete networks (**Figure 4B,C**). Finally at V5, the original subnetworks from V1 (F, G and I) had consolidated into a single large upregulated co-expression module (N, **Figure 4B,C**), which included re-wiring with the now upregulated interferon-associated genes which were originally within the (then quiescent) module A at V1 (**Figure 4B,C**).

To visualise these progressive changes in network structure during the course of SCIT we plotted the top 800 network edges/connectivity patterns from *WGCNA* employing Cytoscape, focusing initially on the components of the key modules A/F/G/I from V1. **Figure 5/Figure E10** illustrates the major structural changes that occurred in the network between V1 to V5, notably the merging of the initially discrete Th2-, IL2-, and Interferon-signalling modules into a single interconnected coexpression module by the end of 2yrs treatment. As shown this transition appears to involve two stages, with initial coalescing of the Th2-and IL2-signalling modules evident by the end of SCIT updosing(V2), while final merging with the Interferon module was not observed until the second year of SCIT(V5). Of note, repetition of these analyses employing other network parameters (softPower 12,13, and minimum module size=20), resulted in similar conclusions regarding network rewiring (data not shown).

**Figure 5.**
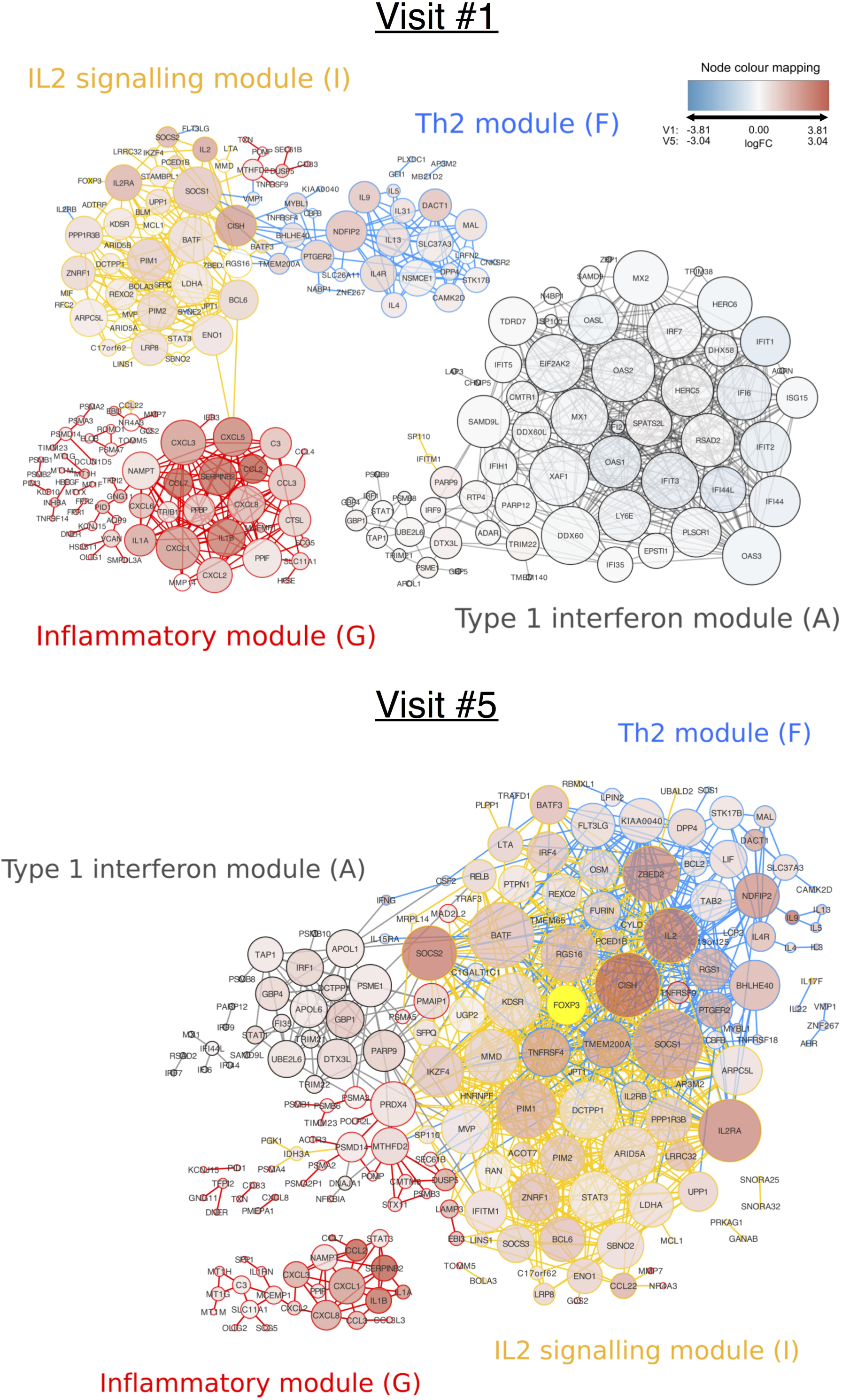
Reconstruction of the network wiring diagram at V1/V5 reveals changes in network connectivity. Co-expression networks were constructed employing *WGCNA* and visualised in Cytosape (n=25, V1; n=24, V5). The top-800 network edges/pairwise connections were utilised for network reconstruction. We repeated this analysis with a different number of edges (e.g. 300, 500, 800, 1000, 1500) and the data were unchanged. The node size reflects the number of connections per gene. Node colour spectrum in key (red-blue) designates positive-to-negative log2FC; note module A is not upregulated until V5.

We also calculated network summary statistics to identify hubs and bottleneck nodes^21^ that are thought to play central roles in network stabilisation. With respect to the key V5 module N, 10 of the top 20 hub/bottleneck genes identified via Cytoscape based on the betweenness centrality and degree metrics^21^ were associated with IL2/FOXP3-signalling and 4 with Type1 IFN-signalling (**Table E8**), consistent with their likely contributions to eventual consolidation of the SCIT-induced low responder phenotype.

As noted in Methods 18 of the 25 HDM allergic subjects employed here were also grass responsive, and received grass SCIT in parallel with HDM treatment. We repeated these analyses on HDM associated responses after removal of the 7 subjects who did not receive grass SCIT (**Figure E11**), and the overall HDM-associated response pattern (in particular the module consolidation at V5; **Figure E12**) was essentially unchanged.

## DISCUSSION

Successful SCIT is typically a biphasic process, in which symptom improvement commences in responsive subjects within a few months and peaks within the first treatment year, however “consolidation” of the clinical effects of SCIT requires a 2^nd^ (sometimes 3^rd^) year of maintenance treatment to prevent slow reversion to pre-treatment allergen-responder status^22,23^. Current understanding of the mechanism(s) underlying desensitisation derive principally from studies on the early phase of SCIT, which suggest important roles for T-regulatory cells, allergen-specific IgG4, and a broad range of other negative control mechanisms^1,2,4^. How these mechanisms interact to control allergen-induced responses in this early phase of SCIT is incompletely understood, and it is not known whether the same range of mechanisms are responsible for eventual stabilisation of the SCIT-induced low-allergen-responder phenotype, or alternatively whether additional control mechanisms are induced/recruited via more prolonged treatment. In this regard it is pertinent to note that (as in **Figure E7**) specific IgE production is maintained or transiently boosted during the early period of SCIT, and declines only slowly with continuing therapy, even in subjects with markedly reduced symptoms^1,2,24^, inferring that Th2-polarised immunological memory against the target allergen is not completely abrogated by SCIT, but instead its downstream immunoinflammatory effects are attenuated.

Our principal aim in this study was to test whether unbiased systems-level analyses of HDM-specific Th-memory-associated response profiles spanning the early and later(consolidation) phases of SCIT, could provide mechanistic insight into this biphasic process. Our study yielded two sets of complimentary findings. In untreated/highly symptomatic subjects, the HDM-induced Th-memory-associated signature is enriched for multiple genes that map to pathways associated with transcriptional activation and cytokine/cytokine-receptor interactions, and the molecular drivers of this response are dominated by hallmark Th1/Th2-and inflammation-associated genes (**Figure 2A-C**) which in turn are correlated with symptom scores (**Figure 2D-F**). This correlation was lost by the end of SCIT updosing (at 3.5mths), but analysis of the accompanying “early post-treatment” HDM-specific DEG signature revealed minimal accompanying changes in underlying pathway/upstream driver profiles, and moreover this pattern did not change substantially throughout the initial course of maintenance therapy to 12mths(V4; **Figure 3B,C**), despite a further decline in symptom scores(**Figure 1**). Continuation of maintenance therapy for a second 12mth period resulted in stabilisation of symptom scores by 24mths/V5 at 51%(mean) below baseline(**Figure 1**), and the appearance in the HDM-specific response profile of upregulated IFN-signalling (**Figure 3B**; **Table I**) and Type1-IFN-associated transcriptional regulators (particularly STAT1 and IRF7; **Figure 3C**).

Additional gene co-expression analyses provide a holistic view of the overall HDM-specific Th-memory-associated response which is key to interpretation of these findings. As illustrated in **Figures 4/5/E10**, the use of *WCGNA* enables construction of complex networks comprising functional modules, each encompassing genes which are highly correlated with each other on the basis of co-expression; exemplars in **Figure 5** are modules designated respectively “Th2” which includes the core IL4/IL4R/IL5/IL9/IL13 gene signature, and “IL2-signalling” encompassing IL2/IL2R/FOXP3 and related genes. Tracking the connectivity patterns between these and other functionally coherent co-expression modules within the network across the time-course of SCIT suggests that the influence of therapy on the overall allergen-induced DEG response extends beyond quantitative effects on transcription, and importantly includes coordination/synchronisation of the expression of key sets of genes within the Th-memory-associated response.

Thus at V1(pre-treatment) at which time prominent drivers of symptoms include Th2 effectors exemplified by IL4/IL5 and archetypal inflammatory mediators such as TNF/IL1B/1L18, the network modules associated with Th2-, inflammation-, and IL2/FOXP3-signalling were isolated in the network topology, suggesting they operate relatively independently (modules F,G,I in **Figure 4B**), resulting in a low-level of coordination between the activity of inflammatory and regulatory networks in highly symptomatic subjects. However overall network topology is radically altered during the updosing phase of SCIT such that by V2 the Th2 and IL2/FOXP3 modules together with much of the inflammation subnetwork have coalesced into a single entity (module J/**Figure 4B**), suggesting enhanced coordination of expression of symptom-associated and regulatory genes/pathways, and this coincides with initial symptom decline and termination of the nexus between symptom scores and the overall Th-memory-induced DEG signature. The early SCIT-induced merging of these subnetworks persists in the face of further changes in network topology as therapy continues, culminating during the 2^nd^ year of treatment with additional merging by V5 with the now upregulated IFN-signalling subnetwork, to form the single highly integrated module N (**Figure 4B**) encompassing coordinated Th2-, IL2/FOXP3-, inflammation-, and IFN-associated signalling. In contrast, modules B, C and D encompassing the key “infrastructural” activities (antigen processing/presentation, TCR activation, translation) which underpin the overall Th-memory-associated response, remain as distinct entities throughout.

We performed additional analyses to visualise changes in network wiring across time, with particular emphasis on gene-gene connections in network module N/V5 which encompasses the main inflammatory effector/regulatory pathways following 2yrs SCIT (**Figure 5)**. As noted in Results, subsidiary analyses (**Table E9**) highlighted the prominence of genes associated with IL2/FOXP3-signalling in stabilising this network and hence consolidation of the clinical effects of SCIT. However additional findings in **Table E9** together with **Table I/Figures 4,5** suggest that other factors related to network topology are also involved. In particular, the upregulation during the 2^nd^ treatment year of the initially quiescent Type1-IFN module and its “wiring” into the aeroallergen-specific Th-memory-associated co-expression network at V5, may also play a key role in this stabilisation process.

In this regard, previous studies have identified Type1-IFN-associated DEG signatures in both PBMC-and sputum-derived cellular response profiles in a variety of settings relevant to asthma-related immunophenotypes^24,25^, with the perceived role of IFNs generally interpreted in the context of host responses to microbial triggers of acute exacerbations. However, we^25^ and others have also shown^26,27^ that Type1-IFNs negatively control many of the downstream pro-inflammatory effects of Th2 cytokines, including the IL4/IL13-induced stimulation of expression of receptors governing the endocytic, trafficking, and IgE-binding properties of myeloid cells^25^. It is thus conceivable that this late-stage rewiring of the network during prolonged maintenance SCIT, resulting in the introduction of coordinated Type1-IFN signalling into the aeroallergen-specific Th-memory-associated response profile, may provide a last-line-of-defence to stabilise immunotherapy via targeting key functions of activated Th2-memory cells that evade control via T-regulatory cells and related mechanisms. Given that the primary source of Type1-IFNs are myeloid cells (particularly DC), the findings above suggest that some of the long-term effects of SCIT may include modulation of the capacity of allergen-specific-Th-memory-cells to chemoattract/cluster-with/stimulate myeloid cells that are present within the microenvironment of the Th-memory cells during their interaction with aeroallergen. There are several earlier studies reporting SIT-associated effects on myeloid functions, including Type1-IFN response capacity of DC^28-30^. It is also relevant to note parallel findings from our earlier studies^31^ contrasting the allergen-specific Th2-memory-associated co-expression network in HDM-sensitised atopic children with its counterpart underlying childhood responses to the highly Th2-polarising Diphtheria-tetanus-acellular-pertussis(DTaP) vaccine, the administration of which does not provoke Th2-associated allergic inflammation despite frequent stimulation of high levels of vaccine-antigen-specific IgE^32^. Mirroring the post SCIT-findings above, the clinically benign Th2-polarised DTaP-specific Th-memory-associated response also included upregulation of a prominent Type1-IFN-associated gene subnetwork, which was not observed in the corresponding allergen-specific network response^31^.

This exploratory study has limitations including sample size, lack of a placebo control group, and non-availability of data on Th-memory-associated profiles in Yr3 during SCIT, which require addressing in follow-ups. Importantly, the same network analytic approach needs to be applied to other immunotherapeutic treatments as SCIT may not be representative of all forms of desensitisation. Notwithstanding these limitations, the study for the first time introduces the concept that remodelling of overall allergen-specific Th-memory-associated response network topology, as opposed to solely quantitative effects on expression of regulatory/effector genes within the response network, may be central to the efficacy of allergen-specific immunotherapeutic treatment modalities. Rapidly emerging single-cell RNA-Seq technology provides an avenue for elucidation of these network interactions with higher precision than was possible in this study, and their application may open up new possibilities for therapeutic design relevant to allergy control.

## Abbreviations

(SCIT): Subcutaneous immunotherapy,
(SIT): specific immunotherapy,
(PBMC): peripheral blood mononuclear cells,
(WGCNA): weighted gene coexpression network analysis,
(DEG): differentially expressed genes,
(URA): upstream regulator analysis,
(LPS): bacterial lipopolysaccharide.

