## Supplementary Tables for "Rewiring of mite allergen-specific Th-memory-associated gene networks during immunotherapy"

### Supplementary Table E1. Subject characteristics at recruitment.

| | |
|---|---|
| **Subject characteristics (n=25)** | |
| Age in years (median/range) | 28.9 (19-56.8) |
| **<u>% of population</u>** | |
| Female | 80 |
| Perennial rhinitis | 100 |
| Asthma | 26.9 |
| Allergic conjunctivitis | 11.5 |
| Current wheeze | 56 |
| Eczema | 3.8 |
| **<u>Sensitizations (%)</u>** | |
| HDM | 100 |
| Grasses | 72 |
| HDM + Grasses | 72 |
| Cat | 26.9 |
| Cockroach | 11.5 |
| Mold | 3.8 |
| Dog | 3.8 |
| **<u>Medication (daily, %)</u>** | |
| Inhaled corticosteroids | 15.4 |
| Inhaled corticosteroids + antihistamine | 53.8 |
| Antihistamine | 19.2 |
| Inhaled corticosteroids + LABA | 3.8 |
| Inhaled corticosteroids + LABA + antihistamine | 7.7 |

**Supplementary Table E2. Top 100 differentially expressed genes (up/down) in CD4+ T cells prior to SCIT (V1).**

| ProbeID | EntrezID | Symbol | logFC | AveExpr | t | P.Value | adj.P.Val | B |
|---|---|---|---|---|---|---|---|---|
| 6347_at | 6347 | CCL2 | 3.62 | 7.76 | 11.58 | 2.08E-22 | 1.18E-20 | 40.30 |
| 6354_at | 6354 | CCL7 | 3.37 | 6.62 | 11.96 | 2.11E-23 | 1.32E-21 | 42.57 |
| 5055_at | 5055 | SERPINB2 | 3.24 | 7.42 | 12.15 | 6.40E-24 | 4.45E-22 | 43.75 |
| 3553_at | 3553 | IL1B | 3.00 | 8.34 | 10.60 | 7.82E-20 | 2.97E-18 | 34.43 |
| 6374_at | 6374 | CXCL5 | 2.75 | 10.79 | 11.70 | 9.83E-23 | 5.73E-21 | 41.05 |
| 2919_at | 2919 | CXCL1 | 2.26 | 7.97 | 10.16 | 1.15E-18 | 3.45E-17 | 31.77 |
| 2921_at | 2921 | CXCL3 | 2.26 | 6.43 | 10.50 | 1.50E-19 | 5.11E-18 | 33.79 |
| 1154_at | 1154 | CISH | 2.10 | 7.68 | 15.19 | 7.66E-32 | 3.12E-29 | 61.82 |
| 3552_at | 3552 | IL1A | 2.06 | 5.66 | 8.32 | 5.56E-14 | 7.25E-13 | 21.11 |
| 3558_at | 3558 | IL2 | 2.04 | 4.57 | 12.33 | 2.13E-24 | 1.55E-22 | 44.84 |
| 2791_at | 2791 | GNG11 | 1.70 | 5.78 | 9.30 | 1.89E-16 | 3.95E-15 | 26.73 |
| 4316_at | 4316 | MMP7 | 1.66 | 5.60 | 9.02 | 1.01E-15 | 1.82E-14 | 25.07 |
| 3559_at | 3559 | IL2RA | 1.65 | 6.83 | 14.84 | 6.05E-31 | 1.57E-28 | 59.78 |
| 4495_at | 4495 | MT1G | 1.63 | 7.62 | 8.10 | 1.97E-13 | 2.32E-12 | 19.87 |
| 3576_at | 3576 | CXCL8 | 1.59 | 9.94 | 8.17 | 1.37E-13 | 1.68E-12 | 20.22 |
| 6372_at | 6372 | CXCL6 | 1.58 | 3.96 | 6.86 | 1.78E-10 | 1.32E-09 | 13.17 |
| 8835_at | 8835 | SOCS2 | 1.56 | 5.65 | 10.86 | 1.65E-20 | 6.73E-19 | 35.97 |
| 55022_at | 55022 | PID1 | 1.48 | 6.03 | 8.19 | 1.20E-13 | 1.48E-12 | 20.36 |
| 54602_at | 54602 | NDFIP2 | 1.43 | 4.93 | 8.55 | 1.50E-14 | 2.29E-13 | 22.41 |
| 718_at | 718 | C3 | 1.38 | 5.54 | 9.33 | 1.57E-16 | 3.33E-15 | 26.91 |
| 4496_at | 4496 | MT1H | 1.35 | 7.64 | 7.64 | 2.60E-12 | 2.53E-11 | 17.33 |
| 6364_at | 6364 | CCL20 | 1.33 | 6.61 | 8.68 | 7.32E-15 | 1.17E-13 | 23.11 |
| 6348_at | 6348 | CCL3 | 1.32 | 7.63 | 6.16 | 6.76E-09 | 4.03E-08 | 9.61 |
| 79413_at | 79413 | ZBED2 | 1.32 | 4.70 | 10.73 | 3.59E-20 | 1.43E-18 | 35.20 |
| 51339_at | 51339 | DACT1 | 1.31 | 6.09 | 7.17 | 3.41E-11 | 2.76E-10 | 14.79 |
| 3578_at | 3578 | IL9 | 1.31 | 4.31 | 4.47 | 1.59E-05 | 5.63E-05 | 2.09 |
| 2920_at | 2920 | CXCL2 | 1.29 | 8.02 | 6.89 | 1.58E-10 | 1.18E-09 | 13.29 |
| 7980_at | 7980 | TFPI2 | 1.28 | 4.41 | 8.14 | 1.62E-13 | 1.97E-12 | 20.06 |
| 6367_at | 6367 | CCL22 | 1.26 | 9.06 | 10.68 | 5.00E-20 | 1.94E-18 | 34.87 |
| 8651_at | 8651 | SOCS1 | 1.22 | 6.17 | 13.59 | 1.04E-27 | 1.26E-25 | 52.40 |
| 5473_at | 5473 | PPBP | 1.16 | 12.17 | 6.49 | 1.25E-09 | 8.24E-09 | 11.26 |
| 199675_at | 199675 | MCEMP1 | 1.11 | 6.65 | 7.62 | 2.89E-12 | 2.77E-11 | 17.22 |
| 3002_at | 3002 | GZMB | 1.09 | 6.84 | 5.67 | 7.27E-08 | 3.74E-07 | 7.30 |
| 170691_at | 170691 | ADAMTS17 | 1.09 | 5.21 | 13.45 | 2.46E-27 | 2.59E-25 | 51.54 |
| 84830_at | 84830 | ADTRP | 1.08 | 6.19 | 13.54 | 1.46E-27 | 1.70E-25 | 52.06 |
| 1462_at | 1462 | VCAN | 1.05 | 4.05 | 7.45 | 7.34E-12 | 6.62E-11 | 16.30 |
| 6447_at | 6447 | SCG5 | 1.05 | 5.08 | 7.00 | 8.69E-11 | 6.74E-10 | 13.88 |
| 114801_at | 114801 | TMEM200A | 1.03 | 4.99 | 10.16 | 1.13E-18 | 3.42E-17 | 31.79 |
| 5732_at | 5732 | PTGER2 | 1.03 | 8.32 | 9.13 | 5.07E-16 | 9.74E-15 | 25.75 |

| ProbeID | EntrezID | Symbol | logFC | AveExpr | t | P.Value | adj.P.Val | B |
|---|---|---|---|---|---|---|---|---|
| 3557_at | 3557 | IL1RN | 1.02 | 6.94 | 5.65 | 8.33E-08 | 4.23E-07 | 7.17 |
| 5292_at | 5292 | PIM1 | 1.02 | 8.90 | 17.87 | 1.40E-38 | 4.58E-35 | 77.17 |
| 4603_at | 4603 | MYBL1 | 0.98 | 8.36 | 9.27 | 2.26E-16 | 4.67E-15 | 26.55 |
| 705_at | 705 | BYSL | 0.98 | 6.53 | 14.38 | 9.35E-30 | 1.70E-27 | 57.06 |
| 1847_at | 1847 | DUSP5 | 0.96 | 7.01 | 8.47 | 2.35E-14 | 3.44E-13 | 21.96 |
| 3562_at | 3562 | IL3 | 0.95 | 3.28 | 4.63 | 8.01E-06 | 3.01E-05 | 2.75 |
| 10148_at | 10148 | EBI3 | 0.95 | 5.53 | 9.21 | 3.26E-16 | 6.57E-15 | 26.19 |
| 6351_at | 6351 | CCL4 | 0.94 | 9.19 | 5.54 | 1.39E-07 | 6.91E-07 | 6.67 |
| 27074_at | 27074 | LAMP3 | 0.94 | 6.31 | 7.31 | 1.60E-11 | 1.36E-10 | 15.54 |
| 25902_at | 25902 | MTHFD1L | 0.94 | 7.03 | 17.58 | 7.26E-38 | 1.18E-34 | 75.55 |
| 645784_at | 645784 | ANKRD36BP2 | 0.93 | 5.12 | 4.77 | 4.36E-06 | 1.70E-05 | 3.34 |
| 22797_at | 22797 | TFEC | -0.76 | 5.45 | -6.10 | 9.19E-09 | 5.38E-08 | 9.31 |
| 5836_at | 5836 | PYGL | -0.78 | 5.39 | -5.79 | 4.09E-08 | 2.20E-07 | 7.86 |
| 1675_at | 1675 | CFD | -0.79 | 4.75 | -8.58 | 1.26E-14 | 1.95E-13 | 22.58 |
| 3597_at | 3597 | IL13RA1 | -0.80 | 5.44 | -6.34 | 2.70E-09 | 1.69E-08 | 10.51 |
| 7305_at | 7305 | TYROBP | -0.81 | 10.08 | -5.71 | 6.15E-08 | 3.23E-07 | 7.46 |
| 51313_at | 51313 | FAM198B | -0.81 | 3.58 | -9.82 | 8.77E-18 | 2.31E-16 | 29.76 |
| 7850_at | 7850 | IL1R2 | -0.82 | 6.73 | -4.19 | 4.84E-05 | 1.58E-04 | 1.03 |
| 2681_at | 2681 | GGTA1P | -0.82 | 3.58 | -8.60 | 1.13E-14 | 1.77E-13 | 22.68 |
| 4286_at | 4286 | MITF | -0.83 | 6.07 | -7.46 | 7.21E-12 | 6.52E-11 | 16.32 |
| 6614_at | 6614 | SIGLEC1 | -0.83 | 5.61 | -6.01 | 1.43E-08 | 8.23E-08 | 8.88 |
| 54674_at | 54674 | LRRN3 | -0.84 | 10.54 | -6.59 | 7.51E-10 | 5.07E-09 | 11.76 |
| 1536_at | 1536 | CYBB | -0.84 | 7.54 | -5.92 | 2.26E-08 | 1.27E-07 | 8.44 |
| 10288_at | 10288 | LILRB2 | -0.87 | 6.24 | -4.81 | 3.68E-06 | 1.45E-05 | 3.50 |
| 10410_at | 10410 | IFITM3 | -0.88 | 6.59 | -4.94 | 2.10E-06 | 8.58E-06 | 4.04 |
| 928_at | 928 | CD9 | -0.88 | 8.26 | -7.62 | 2.90E-12 | 2.77E-11 | 17.22 |
| 2359_at | 2359 | FPR3 | -0.89 | 6.59 | -6.02 | 1.35E-08 | 7.79E-08 | 8.94 |
| 120892_at | 120892 | LRRK2 | -0.89 | 5.13 | -7.94 | 4.87E-13 | 5.39E-12 | 18.98 |
| 7941_at | 7941 | PLA2G7 | -0.90 | 8.92 | -5.67 | 7.43E-08 | 3.82E-07 | 7.28 |
| 1050_at | 1050 | CEBPA | -0.90 | 6.74 | -7.78 | 1.24E-12 | 1.28E-11 | 18.06 |
| 6252_at | 6252 | RTN1 | -0.90 | 5.83 | -8.78 | 4.02E-15 | 6.72E-14 | 23.71 |
| 3109_at | 3109 | HLA-DMB | -0.91 | 6.84 | -9.02 | 9.85E-16 | 1.80E-14 | 25.10 |
| 84417_at | 84417 | C2orf40 | -0.92 | 6.13 | -10.28 | 5.48E-19 | 1.74E-17 | 32.50 |
| 6369_at | 6369 | CCL24 | -0.96 | 9.65 | -5.30 | 4.15E-07 | 1.91E-06 | 5.61 |
| 2662_at | 2662 | GDF10 | -0.97 | 3.92 | -10.50 | 1.50E-19 | 5.11E-18 | 33.78 |
| 5166_at | 5166 | PDK4 | -0.99 | 4.20 | -9.85 | 7.30E-18 | 1.95E-16 | 29.94 |
| 54209_at | 54209 | TREM2 | -1.00 | 5.13 | -9.25 | 2.49E-16 | 5.09E-15 | 26.45 |
| 3627_at | 3627 | CXCL10 | -1.00 | 5.05 | -3.54 | 5.41E-04 | 1.44E-03 | -1.25 |
| 1823_at | 1823 | DSC1 | -1.02 | 5.50 | -9.49 | 6.18E-17 | 1.39E-15 | 27.83 |
| 6039_at | 6039 | RNASE6 | -1.03 | 6.97 | -7.77 | 1.29E-12 | 1.32E-11 | 18.02 |
| 54_at | 54 | ACP5 | -1.05 | 8.58 | -10.43 | 2.27E-19 | 7.35E-18 | 33.37 |
| 200315_at | 200315 | APOBEC3A | -1.07 | 4.78 | -4.93 | 2.24E-06 | 9.10E-06 | 3.98 |

| ProbeID | EntrezID | Symbol | logFC | AveExpr | t | P.Value | adj.P.Val | B |
|---|---|---|---|---|---|---|---|---|
| 2214_at | 2214 | FCGR3A | -1.08 | 5.99 | -4.63 | 8.01E-06 | 3.01E-05 | 2.75 |
| 26253_at | 26253 | CLEC4E | -1.10 | 6.17 | -8.86 | 2.53E-15 | 4.28E-14 | 24.16 |
| 55016_at | 55016 | MARCH1 | -1.11 | 4.41 | -10.22 | 8.06E-19 | 2.48E-17 | 32.12 |
| 2219_at | 2219 | FCN1 | -1.17 | 5.88 | -6.82 | 2.28E-10 | 1.66E-09 | 12.93 |
| 51311_at | 51311 | TLR8 | -1.17 | 5.23 | -8.13 | 1.66E-13 | 2.01E-12 | 20.04 |
| 10170_at | 10170 | DHRS9 | -1.18 | 5.56 | -10.58 | 9.06E-20 | 3.36E-18 | 34.29 |
| 216_at | 216 | ALDH1A1 | -1.19 | 3.62 | -10.54 | 1.14E-19 | 4.04E-18 | 34.06 |
| 4069_at | 4069 | LYZ | -1.25 | 10.88 | -8.21 | 1.07E-13 | 1.34E-12 | 20.47 |
| 10457_at | 10457 | GPNMB | -1.30 | 6.47 | -7.86 | 7.55E-13 | 8.05E-12 | 18.54 |
| 54504_at | 54504 | CPVL | -1.30 | 3.93 | -8.96 | 1.38E-15 | 2.43E-14 | 24.76 |
| 4332_at | 4332 | MNDA | -1.43 | 5.49 | -7.97 | 4.06E-13 | 4.56E-12 | 19.15 |
| 2167_at | 2167 | FABP4 | -1.48 | 6.90 | -7.22 | 2.62E-11 | 2.14E-10 | 15.05 |
| 4199_at | 4199 | ME1 | -1.58 | 5.39 | -12.55 | 5.64E-25 | 4.38E-23 | 46.16 |
| 3957_at | 3957 | LGALS2 | -1.59 | 5.49 | -5.83 | 3.50E-08 | 1.89E-07 | 8.01 |
| 948_at | 948 | CD36 | -1.61 | 5.46 | -10.60 | 8.18E-20 | 3.07E-18 | 34.39 |
| 10875_at | 10875 | FGL2 | -1.61 | 8.18 | -10.34 | 3.91E-19 | 1.25E-17 | 32.84 |
| 7045_at | 7045 | TGFBI | -1.65 | 7.64 | -9.80 | 9.90E-18 | 2.59E-16 | 29.64 |
| 6035_at | 6035 | RNASE1 | -1.86 | 6.30 | -9.10 | 6.03E-16 | 1.14E-14 | 25.58 |
| 79887_at | 79887 | PLBD1 | -2.07 | 6.48 | -14.53 | 3.83E-30 | 7.82E-28 | 57.95 |

Differentially expressed genes were identified with *limma* contrasting HDM versus CTRL in CD4+ T cells prior to SCIT (V1). ProbeID = , EntrezID = , logFC = log base 2 fold change, AveExpr = average log2-expression level for that gene across all the arrays, t = moderated t-statistic, P.value = the associated p-value, adj.P.Val = the Benjamini & Hochberg adjusted P.value for multiple testing, B = the B-statistic is the log-odds that the gene is differentially expressed.

**Supplementary Table E3. Top 10 up-and-down regulated Go annotations at each visit.**

| Up/down | Pathway Name | Pathway Id | Source Name | Adjusted P-value |
|---|---|---|---|---|
| **Visit # 1** | | | | |
| **upregulated Go terms** | poly(A) RNA binding | GO:0044822 | molecular function | 1.25E-26 |
| | immune response | GO:0006955 | biological process | 1.06E-21 |
| | cytosol | GO:0005829 | cellular component | 5.55E-18 |
| | mitochondrion | GO:0005739 | cellular component | 5.43E-16 |
| | gene expression | GO:0010467 | biological process | 2.10E-14 |
| | inflammatory response | GO:0006954 | biological process | 9.03E-14 |
| | extracellular vesicular exosome | GO:0070062 | cellular component | 3.60E-13 |
| | RNA metabolic process | GO:0016070 | biological process | 5.50E-13 |
| | innate immune response | GO:0045087 | biological process | 2.89E-12 |
| | chemokine activity | GO:0008009 | molecular function | 3.37E-12 |
| **downregulated Go terms** | extracellular vesicular exosome | GO:0070062 | cellular component | 4.88E-12 |
| | innate immune response | GO:0045087 | biological process | 8.58E-11 |
| | lysosome | GO:0005764 | cellular component | 8.83E-10 |
| | immune response | GO:0006955 | biological process | 3.05E-09 |
| | lysosomal membrane | GO:0005765 | cellular component | 3.53E-07 |
| | plasma membrane | GO:0005886 | cellular component | 2.10E-06 |
| | external side of plasma membrane | GO:0009897 | cellular component | 1.01E-05 |
| | inflammatory response | GO:0006954 | biological process | 1.14E-05 |
| | cytokine-mediated signaling pathway | GO:0019221 | biological process | 1.33E-05 |
| | signal transduction | GO:0007165 | biological process | 1.90E-05 |
| **Visit # 2** | | | | |
| **upregulated Go terms** | poly(A) RNA binding | GO:0044822 | molecular function | 2.70E-32 |
| | cytosol | GO:0005829 | cellular component | 7.59E-25 |
| | extracellular vesicular exosome | GO:0070062 | cellular component | 3.91E-22 |
| | mitochondrion | GO:0005739 | cellular component | 4.13E-21 |
| | gene expression | GO:0010467 | biological process | 2.95E-19 |
| | immune response | GO:0006955 | biological process | 8.52E-17 |
| | RNA metabolic process | GO:0016070 | biological process | 2.42E-16 |
| | viral process | GO:0016032 | biological process | 5.79E-16 |
| | nucleolus | GO:0005730 | cellular component | 7.62E-13 |
| | regulation of cellular amino acid metabolic process | GO:0006521 | biological process | 7.86E-13 |
| **downregulated Go terms** | innate immune response | GO:0045087 | biological process | 3.01E-12 |
| | extracellular vesicular exosome | GO:0070062 | cellular component | 9.11E-09 |
| | immune response | GO:0006955 | biological process | 3.21E-08 |
| | lysosome | GO:0005764 | cellular component | 1.66E-07 |
| | external side of plasma membrane | GO:0009897 | cellular component | 2.85E-07 |
| | inflammatory response | GO:0006954 | biological process | 3.25E-07 |
| | lysosomal membrane | GO:0005765 | cellular component | 9.74E-06 |
| | cytosol | GO:0005829 | cellular component | 1.58E-05 |
| | signal transduction | GO:0007165 | biological process | 5.30E-05 |

| Up/down | Pathway Name | Pathway Id | Source Name | Adjusted P-value |
|---|---|---|---|---|
| | T cell receptor signaling pathway | GO:0050852 | biological process | 5.32E-05 |
| **Visit # 4** | | | | |
| **upregulated Go terms** | poly(A) RNA binding | GO:0044822 | molecular function | 2.23E-29 |
| | cytosol | GO:0005829 | cellular component | 7.25E-22 |
| | gene expression | GO:0010467 | biological process | 7.39E-21 |
| | mitochondrion | GO:0005739 | cellular component | 2.82E-17 |
| | RNA metabolic process | GO:0016070 | biological process | 3.31E-17 |
| | immune response | GO:0006955 | biological process | 6.32E-17 |
| | extracellular vesicular exosome | GO:0070062 | cellular component | 2.85E-14 |
| | viral process | GO:0016032 | biological process | 3.55E-14 |
| | nucleolus | GO:0005730 | cellular component | 7.46E-13 |
| | mRNA metabolic process | GO:0016071 | biological process | 1.40E-12 |
| **downregulated Go terms** | extracellular vesicular exosome | GO:0070062 | cellular component | 6.80E-15 |
| | innate immune response | GO:0045087 | biological process | 7.13E-15 |
| | immune response | GO:0006955 | biological process | 9.98E-12 |
| | plasma membrane | GO:0005886 | cellular component | 1.37E-11 |
| | lysosome | GO:0005764 | cellular component | 7.50E-11 |
| | external side of plasma membrane | GO:0009897 | cellular component | 1.23E-09 |
| | cytokine-mediated signaling pathway | GO:0019221 | biological process | 1.14E-08 |
| | signal transduction | GO:0007165 | biological process | 7.01E-08 |
| | blood coagulation | GO:0007596 | biological process | 2.44E-07 |
| | inflammatory response | GO:0006954 | biological process | 3.33E-07 |
| **Visit # 5** | | | | |
| **upregulated Go terms** | poly(A) RNA binding | GO:0044822 | molecular function | 3.98E-35 |
| | mitochondrion | GO:0005739 | cellular component | 2.32E-26 |
| | cytosol | GO:0005829 | cellular component | 1.97E-23 |
| | gene expression | GO:0010467 | biological process | 1.56E-19 |
| | extracellular vesicular exosome | GO:0070062 | cellular component | 4.89E-18 |
| | viral process | GO:0016032 | biological process | 5.04E-18 |
| | immune response | GO:0006955 | biological process | 1.22E-14 |
| | RNA metabolic process | GO:0016070 | biological process | 1.40E-14 |
| | regulation of apoptotic process | GO:0042981 | biological process | 1.40E-14 |
| | cytoplasm | GO:0005737 | cellular component | 2.22E-14 |
| **downregulated Go terms** | extracellular vesicular exosome | GO:0070062 | cellular component | 9.50E-20 |
| | innate immune response | GO:0045087 | biological process | 1.94E-13 |
| | lysosome | GO:0005764 | cellular component | 2.75E-13 |
| | plasma membrane | GO:0005886 | cellular component | 1.73E-08 |
| | signal transduction | GO:0007165 | biological process | 1.90E-08 |
| | cytosol | GO:0005829 | cellular component | 2.21E-08 |
| | lysosomal membrane | GO:0005765 | cellular component | 2.34E-08 |
| | blood coagulation | GO:0007596 | biological process | 9.34E-08 |
| | external side of plasma membrane | GO:0009897 | cellular component | 2.84E-07 |
| | leukocyte migration | GO:0050900 | biological process | 7.66E-07 |

**Supplementary Table E4. Association of respiratory symptoms with gene expression prior to SCIT (V1).**

| ProbeID | EntrezID | Symbol | logFC | AveExpr | t | P.Value | adj.P.Val | B |
|---|---|---|---|---|---|---|---|---|
| 3578_at | 3578 | IL9 | 0.11 | 4.06 | 5.86 | 4.32E-07 | 5.83E-04 | 4.94 |
| 3567_at | 3567 | IL5 | 0.07 | 3.33 | 4.64 | 2.73E-05 | 3.82E-03 | 0.85 |
| 386653_at | 386653 | IL31 | 0.07 | 3.70 | 5.51 | 1.46E-06 | 7.93E-04 | 3.74 |
| 54602_at | 54602 | NDFIP2 | 0.06 | 4.88 | 4.79 | 1.67E-05 | 3.59E-03 | 1.33 |
| 3596_at | 3596 | IL13 | 0.05 | 4.59 | 5.10 | 5.87E-06 | 1.90E-03 | 2.36 |
| 3458_at | 3458 | IFNG | 0.05 | 4.45 | 4.59 | 3.30E-05 | 4.14E-03 | 0.67 |
| 3002_at | 3002 | GZMB | 0.05 | 6.61 | 3.52 | 9.64E-04 | 3.50E-02 | -2.60 |
| 1154_at | 1154 | CISH | 0.04 | 7.50 | 3.69 | 5.74E-04 | 2.57E-02 | -2.10 |
| 3566_at | 3566 | IL4R | 0.04 | 7.30 | 3.76 | 4.61E-04 | 2.24E-02 | -1.89 |
| 5996_at | 5996 | RGS1 | 0.04 | 7.94 | 4.70 | 2.27E-05 | 3.59E-03 | 1.03 |
| 387496_at | 387496 | RASL11A | 0.04 | 4.24 | 4.09 | 1.64E-04 | 1.37E-02 | -0.90 |
| 8651_at | 8651 | SOCS1 | 0.04 | 6.05 | 6.04 | 2.23E-07 | 5.83E-04 | 5.60 |
| 7293_at | 7293 | TNFRSF4 | 0.04 | 6.86 | 4.62 | 2.92E-05 | 3.82E-03 | 0.79 |
| 55509_at | 55509 | BATF3 | 0.03 | 5.51 | 4.72 | 2.10E-05 | 3.59E-03 | 1.11 |
| 5732_at | 5732 | PTGER2 | 0.03 | 8.25 | 3.48 | 1.09E-03 | 3.69E-02 | -2.72 |
| 115362_at | 115362 | GBP5 | 0.03 | 8.94 | 3.62 | 7.15E-04 | 2.92E-02 | -2.31 |
| 84255_at | 84255 | SLC37A3 | 0.03 | 6.28 | 5.11 | 5.73E-06 | 1.90E-03 | 2.39 |
| 3642_at | 3642 | INSM1 | 0.03 | 3.43 | 3.73 | 5.10E-04 | 2.41E-02 | -1.99 |
| 4783_at | 4783 | NFIL3 | 0.03 | 6.21 | 4.69 | 2.31E-05 | 3.59E-03 | 1.02 |
| 3559_at | 3559 | IL2RA | 0.03 | 6.80 | 3.75 | 4.82E-04 | 2.32E-02 | -1.94 |
| 817_at | 817 | CAMK2D | 0.03 | 7.35 | 3.82 | 3.83E-04 | 2.05E-02 | -1.71 |
| 2672_at | 2672 | GFI1 | 0.03 | 7.45 | 4.16 | 1.32E-04 | 1.17E-02 | -0.69 |
| 27314_at | 27314 | RAB30 | 0.03 | 5.40 | 5.08 | 6.39E-06 | 1.90E-03 | 2.28 |
| 170691_at | 170691 | ADAMTS17 | 0.02 | 5.24 | 3.87 | 3.32E-04 | 1.86E-02 | -1.58 |
| 94120_at | 94120 | SYTL3 | 0.02 | 6.68 | 3.77 | 4.51E-04 | 2.23E-02 | -1.87 |
| 3565_at | 3565 | IL4 | 0.02 | 3.73 | 3.58 | 7.93E-04 | 3.12E-02 | -2.41 |
| 197370_at | 197370 | NSMCE1 | 0.02 | 9.45 | 4.29 | 8.65E-05 | 9.11E-03 | -0.27 |
| 3662_at | 3662 | IRF4 | 0.02 | 6.36 | 5.36 | 2.44E-06 | 1.14E-03 | 3.23 |
| 64859_at | 64859 | NABP1 | 0.02 | 7.82 | 5.57 | 1.17E-06 | 7.63E-04 | 3.96 |
| 4118_at | 4118 | MAL | 0.02 | 10.93 | 3.59 | 7.79E-04 | 3.10E-02 | -2.40 |
| 115361_at | 115361 | GBP4 | 0.02 | 6.97 | 3.47 | 1.11E-03 | 3.69E-02 | -2.74 |
| 3659_at | 3659 | IRF1 | 0.02 | 9.00 | 3.47 | 1.12E-03 | 3.69E-02 | -2.74 |
| 1803_at | 1803 | DPP4 | 0.02 | 8.90 | 4.02 | 2.05E-04 | 1.49E-02 | -1.11 |
| 7494_at | 7494 | XBP1 | 0.02 | 9.20 | 3.98 | 2.37E-04 | 1.54E-02 | -1.25 |
| 604_at | 604 | BCL6 | 0.02 | 8.05 | 4.43 | 5.49E-05 | 6.17E-03 | 0.17 |
| 6875_at | 6875 | TAF4B | 0.02 | 6.89 | 4.89 | 1.21E-05 | 3.30E-03 | 1.65 |
| 355_at | 355 | FAS | 0.02 | 7.76 | 4.41 | 5.98E-05 | 6.51E-03 | 0.09 |
| 6004_at | 6004 | RGS16 | 0.02 | 4.34 | 3.67 | 6.22E-04 | 2.61E-02 | -2.18 |
| 6361_at | 6361 | CCL17 | 0.02 | 6.26 | 4.11 | 1.55E-04 | 1.34E-02 | -0.84 |

| ProbeID | EntrezID | Symbol | logFC | AveExpr | t | P.Value | adj.P.Val | B |
|---|---|---|---|---|---|---|---|---|
| 83939_at | 83939 | EIF2A | 0.02 | 6.55 | 4.03 | 2.03E-04 | 1.49E-02 | -1.10 |
| 50616_at | 50616 | IL22 | 0.02 | 4.56 | 3.78 | 4.41E-04 | 2.21E-02 | -1.85 |
| 6641_at | 6641 | SNTB1 | 0.02 | 6.34 | 4.21 | 1.13E-04 | 1.06E-02 | -0.54 |
| 494143_at | 494143 | CHAC2 | 0.02 | 6.37 | 3.36 | 1.56E-03 | 4.72E-02 | -3.06 |
| 10906_at | 10906 | TRAFD1 | 0.02 | 7.23 | 5.26 | 3.45E-06 | 1.41E-03 | 2.89 |
| 10797_at | 10797 | MTHFD2 | 0.02 | 8.84 | 4.65 | 2.67E-05 | 3.82E-03 | 0.88 |
| 3628_at | 3628 | INPP1 | 0.02 | 6.92 | 3.73 | 5.16E-04 | 2.41E-02 | -2.00 |
| 9246_at | 9246 | UBE2L6 | 0.02 | 10.17 | 3.40 | 1.37E-03 | 4.25E-02 | -2.93 |
| 8809_at | 8809 | IL18R1 | 0.02 | 5.96 | 3.46 | 1.14E-03 | 3.72E-02 | -2.76 |
| 10538_at | 10538 | BATF | 0.02 | 8.30 | 4.02 | 2.05E-04 | 1.49E-02 | -1.11 |
| 64332_at | 64332 | NFKBIZ | 0.01 | 10.35 | 4.22 | 1.11E-04 | 1.06E-02 | -0.51 |
| 6890_at | 6890 | TAP1 | 0.01 | 10.08 | 3.67 | 6.08E-04 | 2.61E-02 | -2.16 |
| 10308_at | 10308 | ZNF267 | 0.01 | 8.64 | 3.98 | 2.32E-04 | 1.54E-02 | -1.23 |
| 10653_at | 10653 | SPINT2 | 0.01 | 8.68 | 3.99 | 2.29E-04 | 1.54E-02 | -1.22 |
| 5971_at | 5971 | RELB | 0.01 | 7.98 | 3.55 | 8.86E-04 | 3.29E-02 | -2.52 |
| 4049_at | 4049 | LTA | 0.01 | 6.66 | 3.82 | 3.93E-04 | 2.07E-02 | -1.74 |
| 3560_at | 3560 | IL2RB | 0.01 | 9.95 | 3.40 | 1.36E-03 | 4.25E-02 | -2.93 |
| 23102_at | 23102 | TBC1D2B | 0.01 | 6.89 | 4.23 | 1.04E-04 | 1.03E-02 | -0.46 |
| 81671_at | 81671 | VMP1 | 0.01 | 8.73 | 4.01 | 2.15E-04 | 1.53E-02 | -1.16 |
| 23212_at | 23212 | RRS1 | 0.01 | 9.75 | 4.07 | 1.79E-04 | 1.41E-02 | -0.98 |
| 9262_at | 9262 | STK17B | 0.01 | 8.59 | 4.44 | 5.31E-05 | 6.17E-03 | 0.20 |
| 5292_at | 5292 | PIM1 | 0.01 | 8.85 | 3.56 | 8.53E-04 | 3.24E-02 | -2.48 |
| 3702_at | 3702 | ITK | 0.01 | 9.18 | 5.72 | 6.94E-07 | 5.83E-04 | 4.48 |
| 5236_at | 5236 | PGM1 | 0.01 | 7.65 | 4.19 | 1.19E-04 | 1.08E-02 | -0.58 |
| 5906_at | 5906 | RAP1A | 0.01 | 10.27 | 3.50 | 1.03E-03 | 3.69E-02 | -2.67 |
| 8611_at | 8611 | PLPP1 | 0.01 | 7.56 | 4.07 | 1.75E-04 | 1.41E-02 | -0.96 |
| 836_at | 836 | CASP3 | 0.01 | 7.32 | 3.95 | 2.58E-04 | 1.65E-02 | -1.33 |
| 2023_at | 2023 | ENO1 | 0.01 | 12.30 | 3.67 | 6.22E-04 | 2.61E-02 | -2.18 |
| 10531_at | 10531 | PITRM1 | 0.01 | 7.65 | 3.94 | 2.70E-04 | 1.65E-02 | -1.38 |
| 7702_at | 7702 | ZNF143 | 0.01 | 7.63 | 4.57 | 3.44E-05 | 4.16E-03 | 0.63 |
| 1503_at | 1503 | CTPS1 | 0.01 | 6.76 | 3.69 | 5.83E-04 | 2.57E-02 | -2.12 |
| 57559_at | 57559 | STAMBPL1 | 0.01 | 8.70 | 4.26 | 9.68E-05 | 9.88E-03 | -0.38 |
| 6675_at | 6675 | UAP1 | 0.01 | 7.95 | 4.63 | 2.81E-05 | 3.82E-03 | 0.82 |
| 1615_at | 1615 | DARS | 0.01 | 10.15 | 3.87 | 3.34E-04 | 1.86E-02 | -1.58 |
| 6993_at | 6993 | DYNLT1 | 0.01 | 10.32 | 3.66 | 6.23E-04 | 2.61E-02 | -2.18 |
| 29851_at | 29851 | ICOS | 0.01 | 8.95 | 4.80 | 1.63E-05 | 3.59E-03 | 1.36 |
| 8509_at | 8509 | NDST2 | 0.01 | 7.37 | 3.43 | 1.27E-03 | 4.05E-02 | -2.86 |
| 123920_at | 123920 | CMTM3 | 0.01 | 8.87 | 3.88 | 3.23E-04 | 1.86E-02 | -1.55 |
| 54552_at | 54552 | GNL3L | 0.01 | 8.68 | 3.49 | 1.06E-03 | 3.69E-02 | -2.69 |
| 9595_at | 9595 | CYTIP | 0.01 | 10.47 | 3.87 | 3.36E-04 | 1.86E-02 | -1.59 |
| 388962_at | 388962 | BOLA3 | 0.01 | 8.74 | 3.79 | 4.24E-04 | 2.16E-02 | -1.81 |
| 9188_at | 9188 | DDX21 | 0.01 | 10.17 | 3.94 | 2.68E-04 | 1.65E-02 | -1.37 |

| ProbeID | EntrezID | Symbol | logFC | AveExpr | t | P.Value | adj.P.Val | B |
|---|---|---|---|---|---|---|---|---|
| 51602_at | 51602 | NOP58 | 0.00 | 10.33 | 3.48 | 1.09E-03 | 3.69E-02 | -2.71 |
| 55863_at | 55863 | TMEM126B | 0.00 | 10.27 | 3.70 | 5.56E-04 | 2.52E-02 | -2.07 |
| 3608_at | 3608 | ILF2 | 0.00 | 10.44 | 3.61 | 7.38E-04 | 2.97E-02 | -2.34 |
| 6277_at | 6277 | S100A6 | -0.01 | 11.92 | -3.81 | 4.04E-04 | 2.09E-02 | -1.77 |
| 51696_at | 51696 | HECA | -0.01 | 11.37 | -4.71 | 2.20E-05 | 3.59E-03 | 1.07 |
| 83442_at | 83442 | SH3BGRL3 | -0.01 | 10.01 | -3.34 | 1.65E-03 | 4.94E-02 | -3.11 |
| 6281_at | 6281 | S100A10 | -0.01 | 11.43 | -3.37 | 1.50E-03 | 4.59E-02 | -3.02 |
| 6653_at | 6653 | SORL1 | -0.01 | 11.44 | -3.93 | 2.73E-04 | 1.65E-02 | -1.39 |
| 10365_at | 10365 | KLF2 | -0.01 | 11.67 | -3.57 | 8.23E-04 | 3.16E-02 | -2.45 |
| 11329_at | 11329 | STK38 | -0.01 | 11.07 | -3.71 | 5.38E-04 | 2.47E-02 | -2.04 |
| 6095_at | 6095 | RORA | -0.01 | 9.66 | -4.06 | 1.81E-04 | 1.41E-02 | -0.99 |
| 2533_at | 2533 | FYB1 | -0.01 | 10.89 | -3.99 | 2.26E-04 | 1.54E-02 | -1.20 |
| 64968_at | 64968 | MRPS6 | -0.01 | 11.35 | -3.55 | 8.74E-04 | 3.28E-02 | -2.51 |
| 27106_at | 27106 | ARRDC2 | -0.01 | 9.65 | -3.48 | 1.07E-03 | 3.69E-02 | -2.70 |
| 6932_at | 6932 | TCF7 | -0.01 | 8.50 | -3.37 | 1.49E-03 | 4.58E-02 | -3.01 |
| 51411_at | 51411 | BIN2 | -0.01 | 8.56 | -3.43 | 1.24E-03 | 4.02E-02 | -2.84 |
| 79026_at | 79026 | AHNAK | -0.01 | 8.95 | -3.83 | 3.74E-04 | 2.03E-02 | -1.69 |
| 3987_at | 3987 | LIMS1 | -0.01 | 10.26 | -3.65 | 6.48E-04 | 2.68E-02 | -2.22 |
| 28984_at | 28984 | RGCC | -0.01 | 11.66 | -4.72 | 2.10E-05 | 3.59E-03 | 1.11 |
| 1043_at | 1043 | CD52 | -0.01 | 12.61 | -5.71 | 7.14E-07 | 5.83E-04 | 4.45 |
| 9214_at | 9214 | FCMR | -0.01 | 9.65 | -3.58 | 8.12E-04 | 3.16E-02 | -2.44 |
| 85478_at | 85478 | CCDC65 | -0.01 | 7.04 | -3.40 | 1.36E-03 | 4.25E-02 | -2.93 |
| 26119_at | 26119 | LDLRAP1 | -0.01 | 9.40 | -3.47 | 1.10E-03 | 3.69E-02 | -2.73 |
| 1831_at | 1831 | TSC22D3 | -0.01 | 9.15 | -4.72 | 2.14E-05 | 3.59E-03 | 1.09 |
| 6526_at | 6526 | SLC5A3 | -0.01 | 11.33 | -3.87 | 3.35E-04 | 1.86E-02 | -1.59 |
| 120224_at | 120224 | TMEM45B | -0.01 | 6.91 | -3.49 | 1.04E-03 | 3.69E-02 | -2.68 |
| 3399_at | 3399 | ID3 | -0.02 | 9.29 | -4.84 | 1.40E-05 | 3.51E-03 | 1.51 |
| 10457_at | 10457 | GPNMB | -0.04 | 6.81 | -3.54 | 9.08E-04 | 3.33E-02 | -2.54 |

We tested whether allergen-driven T-cell response patterns were associated with respiratory symptoms scores prior to SCIT (V1) employing *limma*. ProbeID = , EntrezID = , logFC = log base 2 fold change, AveExpr = average log2-expression level for that gene across all the arrays, t = moderated t-statistic, P.value = the associated p-value, adj.P.Val = the Benjamini & Hochberg adjusted P.value for multiple testing, B = the B-statistic is the log-odds that the gene is differentially expressed.

**Supplementary Table E5. Top 100 differentially expressed genes (up/down) in CD4+ T cells during SCIT (V2, 3.5mths).**

| ProbeID | EntrezID | Symbol | logFC | AveExpr | t | P.Value | adj.P.Val | B |
|---|---|---|---|---|---|---|---|---|
| 3553_at | 3553 | IL1B | 4.18 | 8.34 | 15.11 | 1.17E-31 | 3.56E-30 | 61.32 |
| 5055_at | 5055 | SERPINB2 | 4.10 | 7.42 | 15.69 | 3.90E-33 | 1.45E-31 | 64.71 |
| 6347_at | 6347 | CCL2 | 4.03 | 7.76 | 13.16 | 1.46E-26 | 2.31E-25 | 49.62 |
| 4316_at | 4316 | MMP7 | 3.54 | 5.60 | 19.68 | 6.67E-43 | 8.37E-41 | 87.12 |
| 3552_at | 3552 | IL1A | 3.37 | 5.66 | 13.92 | 1.47E-28 | 2.86E-27 | 54.20 |
| 6374_at | 6374 | CXCL5 | 3.28 | 10.79 | 14.22 | 2.38E-29 | 4.79E-28 | 56.02 |
| 2919_at | 2919 | CXCL1 | 3.26 | 7.97 | 14.94 | 3.30E-31 | 9.06E-30 | 60.28 |
| 3558_at | 3558 | IL2 | 3.18 | 4.57 | 19.64 | 8.38E-43 | 1.01E-40 | 86.89 |
| 1154_at | 1154 | CISH | 2.98 | 7.68 | 22.07 | 2.29E-48 | 7.49E-46 | 99.64 |
| 2921_at | 2921 | CXCL3 | 2.91 | 6.43 | 13.79 | 3.15E-28 | 5.97E-27 | 53.44 |
| 6354_at | 6354 | CCL7 | 2.89 | 6.62 | 10.47 | 1.70E-19 | 1.31E-18 | 33.42 |
| 8835_at | 8835 | SOCS2 | 2.64 | 5.65 | 18.69 | 1.48E-40 | 1.34E-38 | 81.74 |
| 3578_at | 3578 | IL9 | 2.59 | 4.31 | 8.98 | 1.22E-15 | 6.52E-15 | 24.61 |
| 3576_at | 3576 | CXCL8 | 2.50 | 9.94 | 13.09 | 2.12E-26 | 3.32E-25 | 49.25 |
| 3559_at | 3559 | IL2RA | 2.35 | 6.83 | 21.57 | 2.98E-47 | 8.10E-45 | 97.09 |
| 54602_at | 54602 | NDFIP2 | 2.34 | 4.93 | 14.31 | 1.42E-29 | 2.98E-28 | 56.53 |
| 6372_at | 6372 | CXCL6 | 2.26 | 3.96 | 10.04 | 2.27E-18 | 1.57E-17 | 30.85 |
| 6364_at | 6364 | CCL20 | 2.22 | 6.61 | 14.80 | 7.53E-31 | 1.92E-29 | 59.46 |
| 4312_at | 4312 | MMP1 | 2.18 | 3.93 | 10.31 | 4.55E-19 | 3.37E-18 | 32.45 |
| 3620_at | 3620 | IDO1 | 2.18 | 7.00 | 11.65 | 1.38E-22 | 1.42E-21 | 40.50 |
| 79413_at | 79413 | ZBED2 | 2.17 | 4.70 | 18.09 | 4.22E-39 | 3.28E-37 | 78.40 |
| 6348_at | 6348 | CCL3 | 2.15 | 7.63 | 10.25 | 6.48E-19 | 4.71E-18 | 32.09 |
| 51339_at | 51339 | DACT1 | 1.99 | 6.09 | 11.08 | 4.39E-21 | 3.82E-20 | 37.06 |
| 3002_at | 3002 | GZMB | 1.96 | 6.84 | 10.40 | 2.69E-19 | 2.02E-18 | 32.97 |
| 3562_at | 3562 | IL3 | 1.90 | 3.28 | 9.46 | 7.56E-17 | 4.51E-16 | 27.37 |
| 8651_at | 8651 | SOCS1 | 1.85 | 6.17 | 21.04 | 4.88E-46 | 9.95E-44 | 94.31 |
| 114801_at | 114801 | TMEM200A | 1.78 | 4.99 | 17.81 | 1.98E-38 | 1.41E-36 | 76.86 |
| 6367_at | 6367 | CCL22 | 1.77 | 9.06 | 15.35 | 2.95E-32 | 9.72E-31 | 62.69 |
| 7293_at | 7293 | TNFRSF4 | 1.75 | 7.04 | 15.75 | 2.74E-33 | 1.08E-31 | 65.06 |
| 2920_at | 2920 | CXCL2 | 1.70 | 8.02 | 9.29 | 2.02E-16 | 1.16E-15 | 26.39 |
| 10148_at | 10148 | EBI3 | 1.68 | 5.53 | 16.68 | 1.27E-35 | 6.28E-34 | 70.41 |
| 1847_at | 1847 | DUSP5 | 1.65 | 7.01 | 14.82 | 6.61E-31 | 1.71E-29 | 59.59 |
| 4495_at | 4495 | MT1G | 1.61 | 7.62 | 8.20 | 1.16E-13 | 5.32E-13 | 20.09 |
| 6363_at | 6363 | CCL19 | 1.60 | 4.18 | 12.18 | 5.40E-24 | 6.30E-23 | 43.73 |
| 5732_at | 5732 | PTGER2 | 1.59 | 8.32 | 14.38 | 8.97E-30 | 1.98E-28 | 56.99 |
| 27074_at | 27074 | LAMP3 | 1.59 | 6.31 | 12.60 | 4.23E-25 | 5.57E-24 | 46.27 |
| 5996_at | 5996 | RGS1 | 1.56 | 8.12 | 14.38 | 8.96E-30 | 1.98E-28 | 56.99 |
| 112744_at | 112744 | IL17F | 1.53 | 4.61 | 10.50 | 1.47E-19 | 1.14E-18 | 33.57 |
| 705_at | 705 | BYSL | 1.53 | 6.53 | 22.78 | 6.33E-50 | 2.58E-47 | 103.22 |

| ProbeID | EntrezID | Symbol | logFC | AveExpr | t | P.Value | adj.P.Val | B |
|---|---|---|---|---|---|---|---|---|
| 6346_at | 6346 | CCL1 | 1.45 | 5.36 | 11.14 | 2.99E-21 | 2.70E-20 | 37.44 |
| 2791_at | 2791 | GNG11 | 1.45 | 5.78 | 8.09 | 2.16E-13 | 9.59E-13 | 19.48 |
| 50810_at | 50810 | HDGFL3 | 1.44 | 6.47 | 13.50 | 1.78E-27 | 3.17E-26 | 51.72 |
| 8553_at | 8553 | BHLHE40 | 1.43 | 7.04 | 16.62 | 1.73E-35 | 8.30E-34 | 70.11 |
| 3566_at | 3566 | IL4R | 1.42 | 7.40 | 12.54 | 5.99E-25 | 7.76E-24 | 45.92 |
| 10409_at | 10409 | BASP1 | 1.42 | 7.65 | 12.85 | 9.23E-26 | 1.30E-24 | 47.78 |
| 3604_at | 3604 | TNFRSF9 | 1.40 | 5.40 | 14.35 | 1.09E-29 | 2.38E-28 | 56.79 |
| 4603_at | 4603 | MYBL1 | 1.40 | 8.36 | 13.49 | 1.97E-27 | 3.43E-26 | 51.62 |
| 5292_at | 5292 | PIM1 | 1.39 | 8.90 | 25.00 | 1.31E-54 | 2.13E-51 | 113.94 |
| 414062_at | 414062 | CCL3L3 | 1.39 | 7.18 | 8.21 | 1.04E-13 | 4.82E-13 | 20.20 |
| 6004_at | 6004 | RGS16 | 1.38 | 4.44 | 19.51 | 1.66E-42 | 1.94E-40 | 86.21 |
| 9332_at | 9332 | CD163 | -1.09 | 4.99 | -4.36 | 2.46E-05 | 5.53E-05 | 1.32 |
| 3434_at | 3434 | IFIT1 | -1.12 | 8.38 | -5.28 | 4.65E-07 | 1.25E-06 | 5.15 |
| 10437_at | 10437 | IFI30 | -1.14 | 11.13 | -8.69 | 6.69E-15 | 3.37E-14 | 22.92 |
| 7850_at | 7850 | IL1R2 | -1.14 | 6.73 | -5.93 | 2.05E-08 | 6.26E-08 | 8.20 |
| 5836_at | 5836 | PYGL | -1.15 | 5.39 | -8.71 | 5.95E-15 | 3.01E-14 | 23.04 |
| 200315_at | 200315 | APOBEC3A | -1.16 | 4.78 | -5.46 | 2.03E-07 | 5.64E-07 | 5.96 |
| 50856_at | 50856 | CLEC4A | -1.17 | 7.81 | -9.93 | 4.38E-18 | 2.95E-17 | 30.19 |
| 22797_at | 22797 | TFEC | -1.17 | 5.45 | -9.58 | 3.68E-17 | 2.26E-16 | 28.08 |
| 2517_at | 2517 | FUCA1 | -1.17 | 7.69 | -13.95 | 1.23E-28 | 2.40E-27 | 54.38 |
| 55016_at | 55016 | MARCH1 | -1.18 | 4.41 | -11.07 | 4.61E-21 | 4.00E-20 | 37.01 |
| 26509_at | 26509 | MYOF | -1.20 | 6.79 | -9.41 | 9.99E-17 | 5.88E-16 | 27.09 |
| 10170_at | 10170 | DHRS9 | -1.20 | 5.56 | -10.99 | 7.40E-21 | 6.34E-20 | 36.54 |
| 54209_at | 54209 | TREM2 | -1.21 | 5.13 | -11.40 | 6.20E-22 | 6.03E-21 | 39.01 |
| 1378_at | 1378 | CR1 | -1.23 | 5.94 | -9.97 | 3.54E-18 | 2.40E-17 | 30.41 |
| 6252_at | 6252 | RTN1 | -1.23 | 5.83 | -12.21 | 4.62E-24 | 5.42E-23 | 43.89 |
| 26253_at | 26253 | CLEC4E | -1.23 | 6.17 | -10.09 | 1.74E-18 | 1.21E-17 | 31.11 |
| 3109_at | 3109 | HLA-DMB | -1.23 | 6.84 | -12.49 | 8.51E-25 | 1.09E-23 | 45.57 |
| 4286_at | 4286 | MITF | -1.23 | 6.07 | -11.29 | 1.23E-21 | 1.17E-20 | 38.32 |
| 2934_at | 2934 | GSN | -1.28 | 8.83 | -9.56 | 4.05E-17 | 2.48E-16 | 27.99 |
| 54_at | 54 | ACP5 | -1.29 | 8.58 | -12.99 | 4.10E-26 | 6.11E-25 | 48.59 |
| 5166_at | 5166 | PDK4 | -1.32 | 4.20 | -13.50 | 1.86E-27 | 3.29E-26 | 51.67 |
| 54504_at | 54504 | CPVL | -1.39 | 3.93 | -9.78 | 1.12E-17 | 7.22E-17 | 29.27 |
| 10261_at | 10261 | IGSF6 | -1.40 | 7.72 | -9.75 | 1.34E-17 | 8.63E-17 | 29.09 |
| 84417_at | 84417 | C2orf40 | -1.40 | 6.13 | -16.05 | 4.72E-34 | 1.99E-32 | 66.81 |
| 1050_at | 1050 | CEBPA | -1.40 | 6.74 | -12.37 | 1.71E-24 | 2.10E-23 | 44.88 |
| 3627_at | 3627 | CXCL10 | -1.41 | 5.05 | -5.06 | 1.26E-06 | 3.23E-06 | 4.19 |
| 51311_at | 51311 | TLR8 | -1.48 | 5.23 | -10.49 | 1.56E-19 | 1.20E-18 | 33.51 |
| 928_at | 928 | CD9 | -1.52 | 8.26 | -13.48 | 2.08E-27 | 3.60E-26 | 51.56 |
| 7305_at | 7305 | TYROBP | -1.56 | 10.08 | -11.26 | 1.43E-21 | 1.34E-20 | 38.18 |
| 6383_at | 6383 | SDC2 | -1.59 | 8.05 | -10.17 | 1.08E-18 | 7.67E-18 | 31.59 |
| 2219_at | 2219 | FCN1 | -1.76 | 5.88 | -10.47 | 1.76E-19 | 1.35E-18 | 33.39 |

| ProbeID | EntrezID | Symbol | logFC | AveExpr | t | P.Value | adj.P.Val | B |
|---|---|---|---|---|---|---|---|---|
| 7941_at | 7941 | PLA2G7 | -1.79 | 8.92 | -11.54 | 2.62E-22 | 2.64E-21 | 39.86 |
| 10288_at | 10288 | LILRB2 | -1.79 | 6.24 | -10.14 | 1.24E-18 | 8.71E-18 | 31.45 |
| 1536_at | 1536 | CYBB | -1.79 | 7.54 | -12.85 | 9.04E-26 | 1.28E-24 | 47.80 |
| 760_at | 760 | CA2 | -1.83 | 6.49 | -13.57 | 1.19E-27 | 2.16E-26 | 52.11 |
| 2359_at | 2359 | FPR3 | -1.87 | 6.59 | -12.99 | 3.93E-26 | 5.89E-25 | 48.63 |
| 6039_at | 6039 | RNASE6 | -1.88 | 6.97 | -14.51 | 4.25E-30 | 9.90E-29 | 57.73 |
| 2214_at | 2214 | FCGR3A | -1.92 | 5.99 | -8.39 | 3.85E-14 | 1.85E-13 | 21.19 |
| 4199_at | 4199 | ME1 | -1.99 | 5.39 | -16.08 | 4.08E-34 | 1.75E-32 | 66.95 |
| 4332_at | 4332 | MNDA | -2.09 | 5.49 | -11.92 | 2.69E-23 | 2.95E-22 | 42.13 |
| 6369_at | 6369 | CCL24 | -2.12 | 9.65 | -11.94 | 2.37E-23 | 2.64E-22 | 42.25 |
| 4069_at | 4069 | LYZ | -2.22 | 10.88 | -14.93 | 3.43E-31 | 9.33E-30 | 60.24 |
| 948_at | 948 | CD36 | -2.29 | 5.46 | -15.39 | 2.29E-32 | 7.71E-31 | 62.94 |
| 10457_at | 10457 | GPNMB | -2.29 | 6.47 | -14.20 | 2.74E-29 | 5.45E-28 | 55.88 |
| 3957_at | 3957 | LGALS2 | -2.36 | 5.49 | -8.83 | 2.92E-15 | 1.51E-14 | 23.74 |
| 10875_at | 10875 | FGL2 | -2.55 | 8.18 | -16.73 | 9.16E-36 | 4.60E-34 | 70.74 |
| 79887_at | 79887 | PLBD1 | -2.71 | 6.48 | -19.47 | 2.01E-42 | 2.19E-40 | 86.02 |
| 7045_at | 7045 | TGFBI | -2.77 | 7.64 | -16.85 | 4.62E-36 | 2.39E-34 | 71.42 |
| 6035_at | 6035 | RNASE1 | -3.15 | 6.30 | -15.72 | 3.27E-33 | 1.26E-31 | 64.88 |
| 2167_at | 2167 | FABP4 | -3.70 | 6.90 | -18.39 | 7.91E-40 | 6.80E-38 | 80.07 |

Differentially expressed genes were identified with *limma* contrasting HDM versus CTRL in CD4+ T cells during SCIT (V2, 3.5mths). ProbeID = , EntrezID = , logFC = log base 2 fold change, AveExpr = average log2-expression level for that gene across all the arrays, t = moderated t-statistic, P.value = the associated p-value, adj.P.Val = the Benjamini & Hochberg adjusted P.value for multiple testing, B = the B-statistic is the log-odds that the gene is differentially expressed.

**Supplementary Table E6. Top 100 differentially expressed genes (up/down) in CD4+ T cells during SCIT (V4, 12mths).**

| ProbeID | EntrezID | Symbol | logFC | AveExpr | t | P.Value | adj.P.Val | B |
|---|---|---|---|---|---|---|---|---|
| 5055_at | 5055 | SERPINB2 | 3.01 | 7.42 | 11.19 | 2.30E-21 | 1.07E-19 | 37.91 |
| 6347_at | 6347 | CCL2 | 3.01 | 7.76 | 9.52 | 5.06E-17 | 1.04E-15 | 28.01 |
| 6354_at | 6354 | CCL7 | 2.93 | 6.62 | 10.32 | 4.26E-19 | 1.24E-17 | 32.74 |
| 3553_at | 3553 | IL1B | 2.77 | 8.34 | 9.69 | 1.83E-17 | 3.98E-16 | 29.01 |
| 1154_at | 1154 | CISH | 2.23 | 7.68 | 16.00 | 6.28E-34 | 3.42E-31 | 66.59 |
| 3558_at | 3558 | IL2 | 2.18 | 4.57 | 13.05 | 2.80E-26 | 2.19E-24 | 49.12 |
| 6374_at | 6374 | CXCL5 | 2.10 | 10.79 | 8.85 | 2.69E-15 | 4.13E-14 | 24.08 |
| 2921_at | 2921 | CXCL3 | 2.01 | 6.43 | 9.21 | 3.16E-16 | 5.52E-15 | 26.20 |
| 3552_at | 3552 | IL1A | 1.98 | 5.66 | 7.93 | 5.27E-13 | 5.58E-12 | 18.87 |
| 2919_at | 2919 | CXCL1 | 1.94 | 7.97 | 8.62 | 9.94E-15 | 1.37E-13 | 22.79 |
| 8835_at | 8835 | SOCS2 | 1.94 | 5.65 | 13.34 | 4.66E-27 | 5.25E-25 | 50.90 |
| 3559_at | 3559 | IL2RA | 1.77 | 6.83 | 15.82 | 1.84E-33 | 7.50E-31 | 65.52 |
| 4316_at | 4316 | MMP7 | 1.72 | 5.60 | 9.26 | 2.40E-16 | 4.34E-15 | 26.47 |
| 51339_at | 51339 | DACT1 | 1.48 | 6.09 | 8.02 | 3.20E-13 | 3.55E-12 | 19.37 |
| 54602_at | 54602 | NDFIP2 | 1.47 | 4.93 | 8.71 | 5.97E-15 | 8.52E-14 | 23.29 |
| 6364_at | 6364 | CCL20 | 1.46 | 6.61 | 9.43 | 8.71E-17 | 1.69E-15 | 27.47 |
| 2791_at | 2791 | GNG11 | 1.44 | 5.78 | 7.80 | 1.11E-12 | 1.13E-11 | 18.14 |
| 6372_at | 6372 | CXCL6 | 1.41 | 3.96 | 6.10 | 9.10E-09 | 5.19E-08 | 9.29 |
| 79413_at | 79413 | ZBED2 | 1.37 | 4.70 | 11.07 | 4.59E-21 | 2.03E-19 | 37.22 |
| 3576_at | 3576 | CXCL8 | 1.36 | 9.94 | 6.90 | 1.45E-10 | 1.05E-09 | 13.35 |
| 8651_at | 8651 | SOCS1 | 1.31 | 6.17 | 14.42 | 7.44E-30 | 1.16E-27 | 57.29 |
| 6348_at | 6348 | CCL3 | 1.23 | 7.63 | 5.70 | 6.47E-08 | 3.18E-07 | 7.38 |
| 114801_at | 114801 | TMEM200A | 1.21 | 4.99 | 11.79 | 5.70E-23 | 3.58E-21 | 41.57 |
| 3562_at | 3562 | IL3 | 1.18 | 3.28 | 5.69 | 6.59E-08 | 3.22E-07 | 7.37 |
| 718_at | 718 | C3 | 1.16 | 5.54 | 7.77 | 1.25E-12 | 1.26E-11 | 18.02 |
| 4495_at | 4495 | MT1G | 1.16 | 7.62 | 5.69 | 6.66E-08 | 3.25E-07 | 7.35 |
| 3578_at | 3578 | IL9 | 1.15 | 4.31 | 3.88 | 1.61E-04 | 4.67E-04 | -0.14 |
| 4603_at | 4603 | MYBL1 | 1.14 | 8.36 | 10.61 | 7.59E-20 | 2.55E-18 | 34.44 |
| 3002_at | 3002 | GZMB | 1.13 | 6.84 | 5.79 | 4.12E-08 | 2.11E-07 | 7.82 |
| 5732_at | 5732 | PTGER2 | 1.12 | 8.32 | 9.83 | 8.12E-18 | 1.85E-16 | 29.82 |
| 2920_at | 2920 | CXCL2 | 1.12 | 8.02 | 5.91 | 2.26E-08 | 1.20E-07 | 8.41 |
| 7980_at | 7980 | TFPI2 | 1.11 | 4.41 | 6.95 | 1.13E-10 | 8.40E-10 | 13.59 |
| 6367_at | 6367 | CCL22 | 1.10 | 9.06 | 9.20 | 3.49E-16 | 6.05E-15 | 26.10 |
| 705_at | 705 | BYSL | 1.08 | 6.53 | 15.67 | 4.46E-33 | 1.62E-30 | 64.64 |
| 84830_at | 84830 | ADTRP | 1.07 | 6.19 | 13.33 | 5.24E-27 | 5.53E-25 | 50.79 |
| 5292_at | 5292 | PIM1 | 1.05 | 8.90 | 18.35 | 9.86E-40 | 1.61E-36 | 79.82 |
| 4496_at | 4496 | MT1H | 1.03 | 7.64 | 5.76 | 4.74E-08 | 2.41E-07 | 7.69 |
| 91523_at | 91523 | PCED1B | 1.02 | 8.77 | 16.97 | 2.29E-36 | 2.49E-33 | 72.15 |
| 55022_at | 55022 | PID1 | 1.02 | 6.03 | 5.58 | 1.14E-07 | 5.35E-07 | 6.83 |

| ProbeID | EntrezID | Symbol | logFC | AveExpr | t | P.Value | adj.P.Val | B |
|---|---|---|---|---|---|---|---|---|
| 170691_at | 170691 | ADAMTS17 | 1.02 | 5.21 | 12.50 | 7.96E-25 | 5.65E-23 | 45.81 |
| 4312_at | 4312 | MMP1 | 1.01 | 3.93 | 4.62 | 8.32E-06 | 2.92E-05 | 2.68 |
| 3624_at | 3624 | INHBA | 1.00 | 6.01 | 9.28 | 2.16E-16 | 3.99E-15 | 26.57 |
| 6447_at | 6447 | SCG5 | 1.00 | 5.08 | 6.61 | 6.88E-10 | 4.64E-09 | 11.82 |
| 25902_at | 25902 | MTHFD1L | 0.99 | 7.03 | 18.41 | 6.91E-40 | 1.61E-36 | 80.17 |
| 6875_at | 6875 | TAF4B | 0.97 | 6.83 | 14.94 | 3.25E-31 | 6.24E-29 | 60.39 |
| 1839_at | 1839 | HBEGF | 0.95 | 4.19 | 9.85 | 7.45E-18 | 1.71E-16 | 29.90 |
| 3566_at | 3566 | IL4R | 0.93 | 7.40 | 7.92 | 5.38E-13 | 5.68E-12 | 18.85 |
| 1847_at | 1847 | DUSP5 | 0.92 | 7.01 | 8.01 | 3.33E-13 | 3.66E-12 | 19.32 |
| 1462_at | 1462 | VCAN | 0.92 | 4.05 | 6.44 | 1.62E-09 | 1.04E-08 | 10.98 |
| 7293_at | 7293 | TNFRSF4 | 0.89 | 7.04 | 7.74 | 1.48E-12 | 1.47E-11 | 17.86 |
| 1823_at | 1823 | DSC1 | -0.83 | 5.50 | -7.65 | 2.46E-12 | 2.36E-11 | 17.35 |
| 3434_at | 3434 | IFIT1 | -0.83 | 8.38 | -3.82 | 2.00E-04 | 5.71E-04 | -0.35 |
| 7077_at | 7077 | TIMP2 | -0.84 | 7.27 | -8.61 | 1.07E-14 | 1.48E-13 | 22.71 |
| 6614_at | 6614 | SIGLEC1 | -0.85 | 5.61 | -6.05 | 1.16E-08 | 6.43E-08 | 9.06 |
| 8714_at | 8714 | ABCC3 | -0.86 | 5.55 | -7.41 | 9.18E-12 | 8.15E-11 | 16.06 |
| 760_at | 760 | CA2 | -0.86 | 6.49 | -6.16 | 6.72E-09 | 3.90E-08 | 9.59 |
| 84417_at | 84417 | C2orf40 | -0.86 | 6.13 | -9.59 | 3.47E-17 | 7.25E-16 | 28.38 |
| 3597_at | 3597 | IL13RA1 | -0.89 | 5.44 | -7.07 | 5.97E-11 | 4.67E-10 | 14.22 |
| 22797_at | 22797 | TFEC | -0.90 | 5.45 | -7.16 | 3.70E-11 | 2.98E-10 | 14.69 |
| 5836_at | 5836 | PYGL | -0.95 | 5.39 | -6.95 | 1.14E-10 | 8.41E-10 | 13.59 |
| 3109_at | 3109 | HLA-DMB | -0.95 | 6.84 | -9.30 | 1.87E-16 | 3.49E-15 | 26.71 |
| 50856_at | 50856 | CLEC4A | -0.97 | 7.81 | -7.97 | 4.28E-13 | 4.59E-12 | 19.08 |
| 7850_at | 7850 | IL1R2 | -0.97 | 6.73 | -4.91 | 2.41E-06 | 9.10E-06 | 3.88 |
| 120892_at | 120892 | LRRK2 | -0.99 | 5.13 | -8.76 | 4.43E-15 | 6.43E-14 | 23.59 |
| 4286_at | 4286 | MITF | -0.99 | 6.07 | -8.81 | 3.41E-15 | 5.07E-14 | 23.85 |
| 928_at | 928 | CD9 | -0.99 | 8.26 | -8.55 | 1.53E-14 | 2.05E-13 | 22.37 |
| 10410_at | 10410 | IFITM3 | -1.04 | 6.59 | -5.82 | 3.62E-08 | 1.87E-07 | 7.95 |
| 7305_at | 7305 | TYROBP | -1.04 | 10.08 | -7.32 | 1.57E-11 | 1.32E-10 | 15.53 |
| 6252_at | 6252 | RTN1 | -1.06 | 5.83 | -10.26 | 6.06E-19 | 1.73E-17 | 32.39 |
| 1536_at | 1536 | CYBB | -1.08 | 7.54 | -7.53 | 4.77E-12 | 4.40E-11 | 16.70 |
| 2359_at | 2359 | FPR3 | -1.09 | 6.59 | -7.33 | 1.44E-11 | 1.22E-10 | 15.62 |
| 54504_at | 54504 | CPVL | -1.10 | 3.93 | -7.50 | 5.82E-12 | 5.30E-11 | 16.51 |
| 1050_at | 1050 | CEBPA | -1.10 | 6.74 | -9.40 | 1.05E-16 | 2.01E-15 | 27.29 |
| 10288_at | 10288 | LILRB2 | -1.10 | 6.24 | -6.05 | 1.14E-08 | 6.30E-08 | 9.08 |
| 55016_at | 55016 | MARCH1 | -1.10 | 4.41 | -10.06 | 2.04E-18 | 5.28E-17 | 31.19 |
| 6369_at | 6369 | CCL24 | -1.12 | 9.65 | -6.09 | 9.29E-09 | 5.28E-08 | 9.27 |
| 2214_at | 2214 | FCGR3A | -1.12 | 5.99 | -4.75 | 4.78E-06 | 1.73E-05 | 3.22 |
| 216_at | 216 | ALDH1A1 | -1.13 | 3.62 | -9.91 | 4.95E-18 | 1.19E-16 | 30.31 |
| 10170_at | 10170 | DHRS9 | -1.13 | 5.56 | -10.06 | 2.09E-18 | 5.34E-17 | 31.16 |
| 5166_at | 5166 | PDK4 | -1.14 | 4.20 | -11.29 | 1.18E-21 | 5.89E-20 | 38.57 |
| 7941_at | 7941 | PLA2G7 | -1.15 | 8.92 | -7.20 | 2.93E-11 | 2.40E-10 | 14.92 |

| ProbeID | EntrezID | Symbol | logFC | AveExpr | t | P.Value | adj.P.Val | B |
|---|---|---|---|---|---|---|---|---|
| 6039_at | 6039 | RNASE6 | -1.16 | 6.97 | -8.67 | 7.45E-15 | 1.05E-13 | 23.07 |
| 26253_at | 26253 | CLEC4E | -1.16 | 6.17 | -9.25 | 2.54E-16 | 4.53E-15 | 26.41 |
| 54_at | 54 | ACP5 | -1.16 | 8.58 | -11.38 | 6.85E-22 | 3.55E-20 | 39.11 |
| 54209_at | 54209 | TREM2 | -1.18 | 5.13 | -10.76 | 3.10E-20 | 1.11E-18 | 35.33 |
| 51311_at | 51311 | TLR8 | -1.25 | 5.23 | -8.59 | 1.21E-14 | 1.65E-13 | 22.60 |
| 3627_at | 3627 | CXCL10 | -1.27 | 5.05 | -4.44 | 1.76E-05 | 5.85E-05 | 1.96 |
| 200315_at | 200315 | APOBEC3A | -1.29 | 4.78 | -5.88 | 2.65E-08 | 1.39E-07 | 8.25 |
| 4069_at | 4069 | LYZ | -1.45 | 10.88 | -9.46 | 7.25E-17 | 1.43E-15 | 27.65 |
| 4332_at | 4332 | MNDA | -1.46 | 5.49 | -8.06 | 2.49E-13 | 2.84E-12 | 19.61 |
| 2219_at | 2219 | FCN1 | -1.48 | 5.88 | -8.56 | 1.44E-14 | 1.95E-13 | 22.42 |
| 10457_at | 10457 | GPNMB | -1.48 | 6.47 | -8.91 | 1.83E-15 | 2.89E-14 | 24.46 |
| 2167_at | 2167 | FABP4 | -1.52 | 6.90 | -7.32 | 1.51E-11 | 1.27E-10 | 15.57 |
| 948_at | 948 | CD36 | -1.69 | 5.46 | -11.01 | 6.85E-21 | 2.94E-19 | 36.83 |
| 3957_at | 3957 | LGALS2 | -1.73 | 5.49 | -6.27 | 3.79E-09 | 2.29E-08 | 10.15 |
| 4199_at | 4199 | ME1 | -1.76 | 5.39 | -13.80 | 2.99E-28 | 3.75E-26 | 53.63 |
| 7045_at | 7045 | TGFBI | -1.82 | 7.64 | -10.71 | 4.13E-20 | 1.45E-18 | 35.05 |
| 10875_at | 10875 | FGL2 | -1.82 | 8.18 | -11.56 | 2.39E-22 | 1.30E-20 | 40.15 |
| 6035_at | 6035 | RNASE1 | -1.94 | 6.30 | -9.39 | 1.09E-16 | 2.07E-15 | 27.25 |
| 79887_at | 79887 | PLBD1 | -2.23 | 6.48 | -15.53 | 9.82E-33 | 2.91E-30 | 63.86 |

Differentially expressed genes were identified with *limma* contrasting HDM versus CTRL in CD4+ T cells during SCIT (V4, 12 mths). ProbeID = , EntrezID = , logFC = log base 2 fold change, AveExpr = average log2-expression level for that gene across all the arrays, t = moderated t-statistic, P.value = the associated p-value, adj.P.Val = the Benjamini & Hochberg adjusted P.value for multiple testing, B = the B-statistic is the log-odds that the gene is differentially expressed.

**Supplementary Table E7. Top 100 differentially expressed genes (up/down) in CD4+ T cells during SCIT (V5, 24mths).**

| ProbeID | EntrezID | Symbol | logFC | AveExpr | t | P.Value | adj.P.Val | B |
|---|---|---|---|---|---|---|---|---|
| 6374_at | 6374 | CXCL5 | 3.39 | 8.33 | 11.51 | 7.02E-12 | 7.50E-11 | 16.98 |
| 6347_at | 6347 | CCL2 | 3.25 | 5.41 | 12.77 | 6.50E-13 | 1.05E-11 | 19.40 |
| 5055_at | 5055 | SERPINB2 | 3.04 | 5.91 | 10.50 | 5.34E-11 | 4.51E-10 | 14.92 |
| 1154_at | 1154 | CISH | 2.69 | 7.34 | 20.71 | 4.95E-18 | 8.61E-16 | 31.28 |
| 3553_at | 3553 | IL1B | 2.62 | 7.23 | 9.65 | 3.22E-10 | 2.18E-09 | 13.09 |
| 3558_at | 3558 | IL2 | 2.42 | 4.19 | 15.85 | 3.80E-15 | 1.69E-13 | 24.61 |
| 3578_at | 3578 | IL9 | 2.41 | 3.89 | 5.98 | 2.26E-06 | 6.56E-06 | 4.11 |
| 8835_at | 8835 | SOCS2 | 2.39 | 4.70 | 17.80 | 2.21E-16 | 1.58E-14 | 27.47 |
| 6354_at | 6354 | CCL7 | 2.35 | 4.62 | 9.77 | 2.48E-10 | 1.74E-09 | 13.36 |
| 4316_at | 4316 | MMP7 | 2.13 | 4.29 | 10.95 | 2.12E-11 | 1.99E-10 | 15.86 |
| 79413_at | 79413 | ZBED2 | 2.10 | 4.09 | 16.07 | 2.74E-15 | 1.33E-13 | 24.94 |
| 3559_at | 3559 | IL2RA | 2.09 | 5.85 | 25.81 | 1.78E-20 | 2.27E-17 | 36.84 |
| 3552_at | 3552 | IL1A | 2.04 | 4.74 | 8.44 | 4.99E-09 | 2.50E-08 | 10.30 |
| 2919_at | 2919 | CXCL1 | 1.91 | 6.74 | 9.06 | 1.20E-09 | 6.94E-09 | 11.76 |
| 54602_at | 54602 | NDFIP2 | 1.90 | 4.40 | 12.36 | 1.39E-12 | 1.96E-11 | 18.63 |
| 6364_at | 6364 | CCL20 | 1.79 | 5.76 | 15.73 | 4.59E-15 | 2.00E-13 | 24.42 |
| 3002_at | 3002 | GZMB | 1.74 | 6.23 | 7.92 | 1.72E-08 | 7.63E-08 | 9.04 |
| 3562_at | 3562 | IL3 | 1.73 | 2.78 | 6.71 | 3.47E-07 | 1.17E-06 | 6.00 |
| 114801_at | 114801 | TMEM200A | 1.70 | 4.91 | 17.46 | 3.55E-16 | 2.23E-14 | 27.00 |
| 8651_at | 8651 | SOCS1 | 1.69 | 5.77 | 20.43 | 6.98E-18 | 1.11E-15 | 30.93 |
| 2921_at | 2921 | CXCL3 | 1.68 | 4.74 | 9.56 | 3.91E-10 | 2.60E-09 | 12.89 |
| 51339_at | 51339 | DACT1 | 1.66 | 5.45 | 10.41 | 6.37E-11 | 5.25E-10 | 14.74 |
| 3576_at | 3576 | CXCL8 | 1.61 | 8.14 | 9.04 | 1.25E-09 | 7.22E-09 | 11.71 |
| 7293_at | 7293 | TNFRSF4 | 1.58 | 6.75 | 12.73 | 6.98E-13 | 1.12E-11 | 19.33 |
| 1847_at | 1847 | DUSP5 | 1.54 | 6.21 | 13.24 | 2.83E-13 | 5.39E-12 | 20.24 |
| 5732_at | 5732 | PTGER2 | 1.52 | 7.76 | 18.87 | 5.17E-17 | 5.21E-15 | 28.93 |
| 5996_at | 5996 | RGS1 | 1.45 | 7.74 | 16.77 | 9.61E-16 | 5.10E-14 | 25.99 |
| 3620_at | 3620 | IDO1 | 1.42 | 5.76 | 14.21 | 5.26E-14 | 1.36E-12 | 21.95 |
| 6367_at | 6367 | CCL22 | 1.41 | 7.99 | 16.84 | 8.73E-16 | 4.84E-14 | 26.09 |
| 5292_at | 5292 | PIM1 | 1.40 | 8.04 | 28.56 | 1.29E-21 | 4.67E-18 | 39.40 |
| 50810_at | 50810 | HDGFL3 | 1.36 | 5.64 | 17.79 | 2.23E-16 | 1.58E-14 | 27.46 |
| 115362_at | 115362 | GBP5 | 1.33 | 8.56 | 14.51 | 3.24E-14 | 8.98E-13 | 22.44 |
| 112744_at | 112744 | IL17F | 1.32 | 4.21 | 5.51 | 7.86E-06 | 2.06E-05 | 2.86 |
| 3604_at | 3604 | TNFRSF9 | 1.30 | 4.71 | 13.28 | 2.64E-13 | 5.12E-12 | 20.31 |
| 8553_at | 8553 | BHLHE40 | 1.30 | 6.73 | 14.49 | 3.36E-14 | 9.23E-13 | 22.40 |
| 6348_at | 6348 | CCL3 | 1.29 | 6.06 | 5.57 | 6.71E-06 | 1.77E-05 | 3.02 |
| 27074_at | 27074 | LAMP3 | 1.29 | 5.00 | 19.11 | 3.78E-17 | 4.25E-15 | 29.25 |
| 3458_at | 3458 | IFNG | 1.29 | 4.45 | 8.53 | 4.02E-09 | 2.06E-08 | 10.52 |
| 25902_at | 25902 | MTHFD1L | 1.26 | 6.36 | 24.97 | 4.20E-20 | 4.01E-17 | 36.00 |

| ProbeID | EntrezID | Symbol | logFC | AveExpr | t | P.Value | adj.P.Val | B |
|---|---|---|---|---|---|---|---|---|
| 7291_at | 7291 | TWIST1 | 1.25 | 3.19 | 11.35 | 9.56E-12 | 9.83E-11 | 16.67 |
| 6875_at | 6875 | TAF4B | 1.24 | 6.28 | 22.11 | 9.39E-19 | 2.63E-16 | 32.93 |
| 705_at | 705 | BYSL | 1.24 | 6.13 | 20.76 | 4.69E-18 | 8.58E-16 | 31.33 |
| 6004_at | 6004 | RGS16 | 1.24 | 4.03 | 18.96 | 4.57E-17 | 4.99E-15 | 29.06 |
| 4603_at | 4603 | MYBL1 | 1.23 | 7.95 | 18.74 | 6.18E-17 | 5.91E-15 | 28.75 |
| 170691_at | 170691 | ADAMTS17 | 1.20 | 4.91 | 15.27 | 9.37E-15 | 3.35E-13 | 23.69 |
| 3566_at | 3566 | IL4R | 1.20 | 6.80 | 11.02 | 1.86E-11 | 1.78E-10 | 15.99 |
| 3596_at | 3596 | IL13 | 1.17 | 3.87 | 6.09 | 1.72E-06 | 5.08E-06 | 4.39 |
| 4312_at | 4312 | MMP1 | 1.15 | 3.13 | 4.81 | 5.15E-05 | 1.20E-04 | 0.98 |
| 84830_at | 84830 | ADTRP | 1.12 | 6.02 | 19.74 | 1.67E-17 | 2.36E-15 | 30.06 |
| 84937_at | 84937 | ZNRF1 | 1.11 | 6.62 | 19.13 | 3.68E-17 | 4.25E-15 | 29.27 |
| 8714_at | 8714 | ABCC3 | -0.98 | 4.69 | -7.76 | 2.53E-08 | 1.08E-07 | 8.66 |
| 3108_at | 3108 | HLA-DMA | -0.99 | 6.18 | -9.21 | 8.59E-10 | 5.25E-09 | 12.09 |
| 3597_at | 3597 | IL13RA1 | -0.99 | 3.85 | -7.60 | 3.68E-08 | 1.53E-07 | 8.28 |
| 29992_at | 29992 | PILRA | -0.99 | 6.18 | -8.87 | 1.85E-09 | 1.03E-08 | 11.31 |
| 55016_at | 55016 | MARCH1 | -1.01 | 3.11 | -8.12 | 1.05E-08 | 4.86E-08 | 9.55 |
| 2517_at | 2517 | FUCA1 | -1.02 | 6.50 | -8.40 | 5.40E-09 | 2.67E-08 | 10.22 |
| 140807_at | 140807 | KRT72 | -1.04 | 6.09 | -17.52 | 3.26E-16 | 2.15E-14 | 27.08 |
| 120892_at | 120892 | LRRK2 | -1.04 | 3.55 | -7.63 | 3.48E-08 | 1.46E-07 | 8.33 |
| 22797_at | 22797 | TFEC | -1.04 | 3.70 | -7.96 | 1.54E-08 | 6.88E-08 | 9.16 |
| 4286_at | 4286 | MITF | -1.05 | 4.86 | -8.58 | 3.62E-09 | 1.88E-08 | 10.63 |
| 5836_at | 5836 | PYGL | -1.10 | 3.80 | -7.85 | 2.01E-08 | 8.81E-08 | 8.89 |
| 50856_at | 50856 | CLEC4A | -1.12 | 6.34 | -9.91 | 1.86E-10 | 1.34E-09 | 13.65 |
| 928_at | 928 | CD9 | -1.12 | 6.75 | -8.15 | 9.92E-09 | 4.63E-08 | 9.61 |
| 1471_at | 1471 | CST3 | -1.12 | 4.78 | -8.67 | 2.89E-09 | 1.54E-08 | 10.86 |
| 1823_at | 1823 | DSC1 | -1.13 | 4.72 | -14.17 | 5.66E-14 | 1.41E-12 | 21.87 |
| 1378_at | 1378 | CR1 | -1.13 | 5.42 | -15.70 | 4.81E-15 | 2.07E-13 | 24.37 |
| 10437_at | 10437 | IFI30 | -1.13 | 9.57 | -8.84 | 1.96E-09 | 1.09E-08 | 11.26 |
| 6252_at | 6252 | RTN1 | -1.14 | 4.68 | -9.55 | 4.06E-10 | 2.68E-09 | 12.85 |
| 2934_at | 2934 | GSN | -1.16 | 7.25 | -8.02 | 1.35E-08 | 6.11E-08 | 9.29 |
| 1050_at | 1050 | CEBPA | -1.18 | 5.83 | -8.84 | 1.99E-09 | 1.10E-08 | 11.24 |
| 5166_at | 5166 | PDK4 | -1.23 | 3.07 | -9.76 | 2.54E-10 | 1.77E-09 | 13.33 |
| 84417_at | 84417 | C2orf40 | -1.24 | 5.53 | -17.06 | 6.28E-16 | 3.64E-14 | 26.42 |
| 760_at | 760 | CA2 | -1.26 | 4.80 | -7.81 | 2.22E-08 | 9.65E-08 | 8.79 |
| 26253_at | 26253 | CLEC4E | -1.28 | 4.86 | -9.86 | 2.06E-10 | 1.47E-09 | 13.55 |
| 10462_at | 10462 | CLEC10A | -1.29 | 4.00 | -6.90 | 2.10E-07 | 7.34E-07 | 6.51 |
| 2359_at | 2359 | FPR3 | -1.29 | 5.43 | -8.88 | 1.79E-09 | 1.01E-08 | 11.35 |
| 51311_at | 51311 | TLR8 | -1.30 | 4.01 | -8.33 | 6.51E-09 | 3.15E-08 | 10.03 |
| 6383_at | 6383 | SDC2 | -1.31 | 6.07 | -9.10 | 1.10E-09 | 6.45E-09 | 11.84 |
| 54504_at | 54504 | CPVL | -1.36 | 3.09 | -8.01 | 1.37E-08 | 6.20E-08 | 9.28 |
| 2214_at | 2214 | FCGR3A | -1.36 | 4.06 | -7.05 | 1.44E-07 | 5.23E-07 | 6.89 |
| 3109_at | 3109 | HLA-DMB | -1.39 | 5.27 | -10.89 | 2.39E-11 | 2.21E-10 | 15.73 |

| ProbeID | EntrezID | Symbol | logFC | AveExpr | t | P.Value | adj.P.Val | B |
|---|---|---|---|---|---|---|---|---|
| 4199_at | 4199 | ME1 | -1.40 | 3.77 | -8.63 | 3.17E-09 | 1.68E-08 | 10.76 |
| 10261_at | 10261 | IGSF6 | -1.40 | 5.84 | -10.06 | 1.33E-10 | 1.01E-09 | 13.99 |
| 6369_at | 6369 | CCL24 | -1.46 | 7.25 | -8.28 | 7.31E-09 | 3.51E-08 | 9.92 |
| 10288_at | 10288 | LILRB2 | -1.56 | 4.50 | -9.22 | 8.39E-10 | 5.14E-09 | 12.12 |
| 10457_at | 10457 | GPNMB | -1.63 | 4.52 | -11.55 | 6.46E-12 | 7.06E-11 | 17.06 |
| 7305_at | 7305 | TYROBP | -1.64 | 8.42 | -13.33 | 2.39E-13 | 4.72E-12 | 20.42 |
| 1536_at | 1536 | CYBB | -1.70 | 5.64 | -12.86 | 5.53E-13 | 9.27E-12 | 19.56 |
| 7941_at | 7941 | PLA2G7 | -1.73 | 7.07 | -12.37 | 1.36E-12 | 1.93E-11 | 18.65 |
| 6039_at | 6039 | RNASE6 | -1.78 | 5.31 | -11.54 | 6.55E-12 | 7.12E-11 | 17.05 |
| 2219_at | 2219 | FCN1 | -1.83 | 5.10 | -9.53 | 4.17E-10 | 2.74E-09 | 12.83 |
| 948_at | 948 | CD36 | -1.85 | 4.28 | -11.06 | 1.71E-11 | 1.64E-10 | 16.08 |
| 4332_at | 4332 | MNDA | -1.85 | 3.66 | -8.47 | 4.59E-09 | 2.33E-08 | 10.39 |
| 10875_at | 10875 | FGL2 | -1.95 | 6.62 | -10.51 | 5.16E-11 | 4.38E-10 | 14.95 |
| 7045_at | 7045 | TGFBI | -2.08 | 6.21 | -12.14 | 2.10E-12 | 2.71E-11 | 18.21 |
| 3957_at | 3957 | LGALS2 | -2.25 | 4.40 | -9.94 | 1.73E-10 | 1.26E-09 | 13.72 |
| 2167_at | 2167 | FABP4 | -2.30 | 4.61 | -9.39 | 5.75E-10 | 3.64E-09 | 12.50 |
| 4069_at | 4069 | LYZ | -2.34 | 9.29 | -15.34 | 8.50E-15 | 3.12E-13 | 23.79 |
| 6035_at | 6035 | RNASE1 | -2.38 | 5.47 | -10.34 | 7.40E-11 | 6.01E-10 | 14.59 |
| 79887_at | 79887 | PLBD1 | -2.40 | 5.07 | -11.47 | 7.55E-12 | 7.95E-11 | 16.91 |

Differentially expressed genes were identified with *limma* contrasting HDM versus CTRL in CD4+ T cells during SCIT (V5, 24mths). ProbeID = , EntrezID = , logFC = log base 2 fold change, AveExpr = average log2-expression level for that gene across all the arrays, t = moderated t-statistic, P.value = the associated p-value, adj.P.Val = the Benjamini & Hochberg adjusted P.value for multiple testing, B = the B-statistic is the log-odds that the gene is differentially expressed.

### Supplementary Table E8. Cytoscape metrics for the network at V5.

| GeneID | Module assignment V1 | BetweennessCentrality | Degree |
|---|---|---|---|
| PRDX4 | IL2 signalling (I) | 0.1616 | 20 |
| BATF | IL2 signalling (I) | 0.1297 | 43 |
| CISH | Th2 (F) | 0.1243 | 42 |
| NDFIP2 | Th2 (F) | 0.1056 | 20 |
| MTHFD2 | Inflammation (G) | 0.0899 | 11 |
| PSMD14 | Inflammation (G) | 0.0873 | 7 |
| MVP | IL2 signalling (I) | 0.0768 | 26 |
| SOCS1 | IL2 signalling (I) | 0.0725 | 48 |
| DTX3L | Type 1 IFN (A) | 0.0564 | 14 |
| SOCS2 | IL2 signalling (I) | 0.0508 | 31 |
| BHLHE40 | Th2 (F) | 0.0475 | 26 |
| APOL1 | Type 1 IFN (A) | 0.0462 | 12 |
| PSME1 | Type 1 IFN (A) | 0.0458 | 17 |
| PARP9 | Type 1 IFN (A) | 0.0457 | 16 |
| IL2 | IL2 signalling (I) | 0.0445 | 31 |
| PIM1 | IL2 signalling (I) | 0.0442 | 37 |
| PSMA3 | Inflammation (G) | 0.0431 | 5 |
| ARID5A | IL2 signalling (I) | 0.0402 | 32 |
| IL2RA | IL2 signalling (I) | 0.0402 | 40 |
| ZBED2 | IL2 signalling (I) | 0.0374 | 33 |

Network wiring diagrams were constructed utilising the top-weighted 800 gene-gene interactions extracted from the adjacency matrix from the expression data. BetweennessCentrality = genes in the network that have many "shortest paths" going through them, i.e. connector genes. The closeness centrality measure of each gene to another in the network is a number between 0 and 1; Degree = number of connections with other genes in the network.
