## Supplementary Figures E1 - E12 for "Rewiring of mite allergen-specific Th-memory-associated gene networks during immunotherapy"

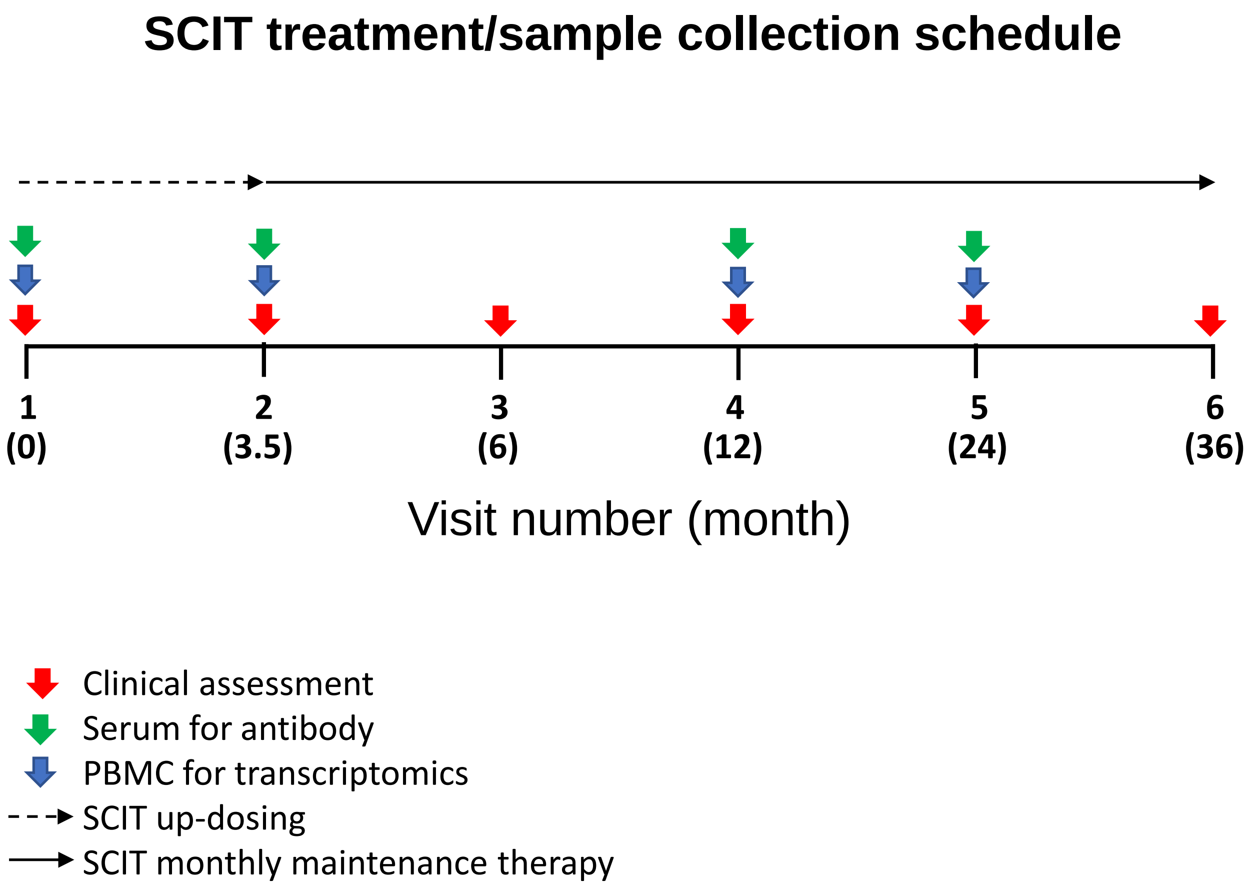


Supplementary Figure E1

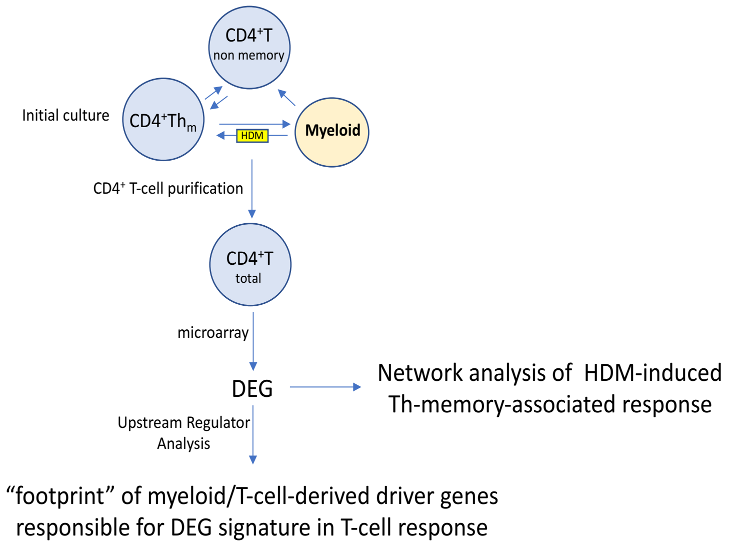


Supplementary Figure E2

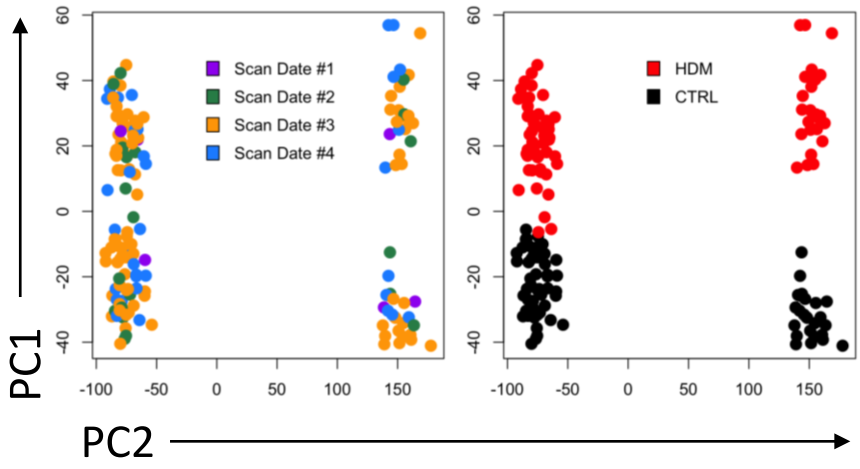


Supplementary Figure E3

PCA plots were utilised to identify batch effects prior to differential expression analysis.

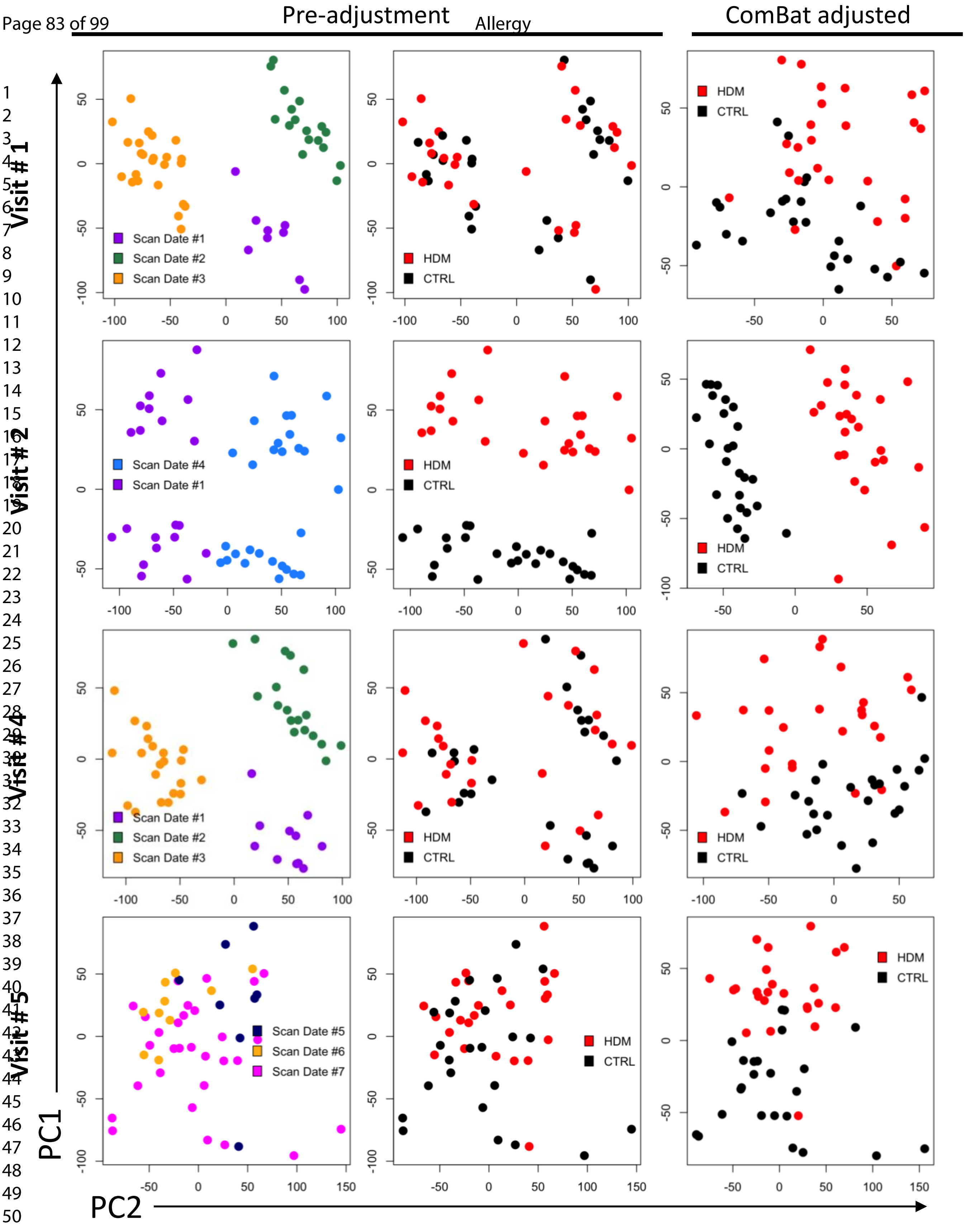


Supplementary Figure E4

PCA plots were utilised to demonstrate batch effect removal with *ComBat* prior to network analysis.

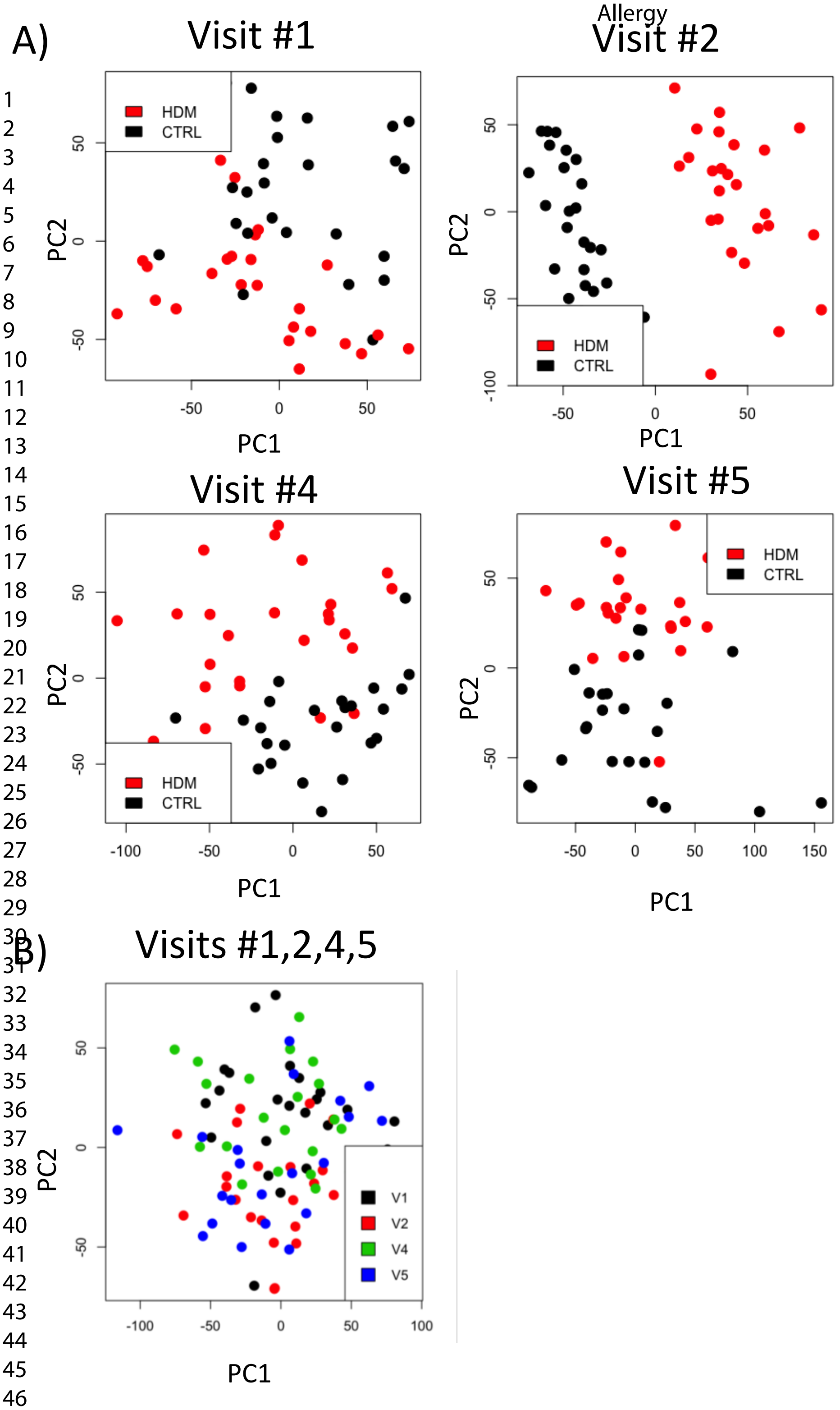


Supplementary Figure E5 - PCA plots of *ComBat* adjusted data at each visit.

A) Principal component analysis for each time point (HDM and medium samples) and B) PCA across all time points (V1, V2, V4 and V5) based on expression log ratios (HDM/ctrl), given that V5 was cultured as a stand-alone experiment.

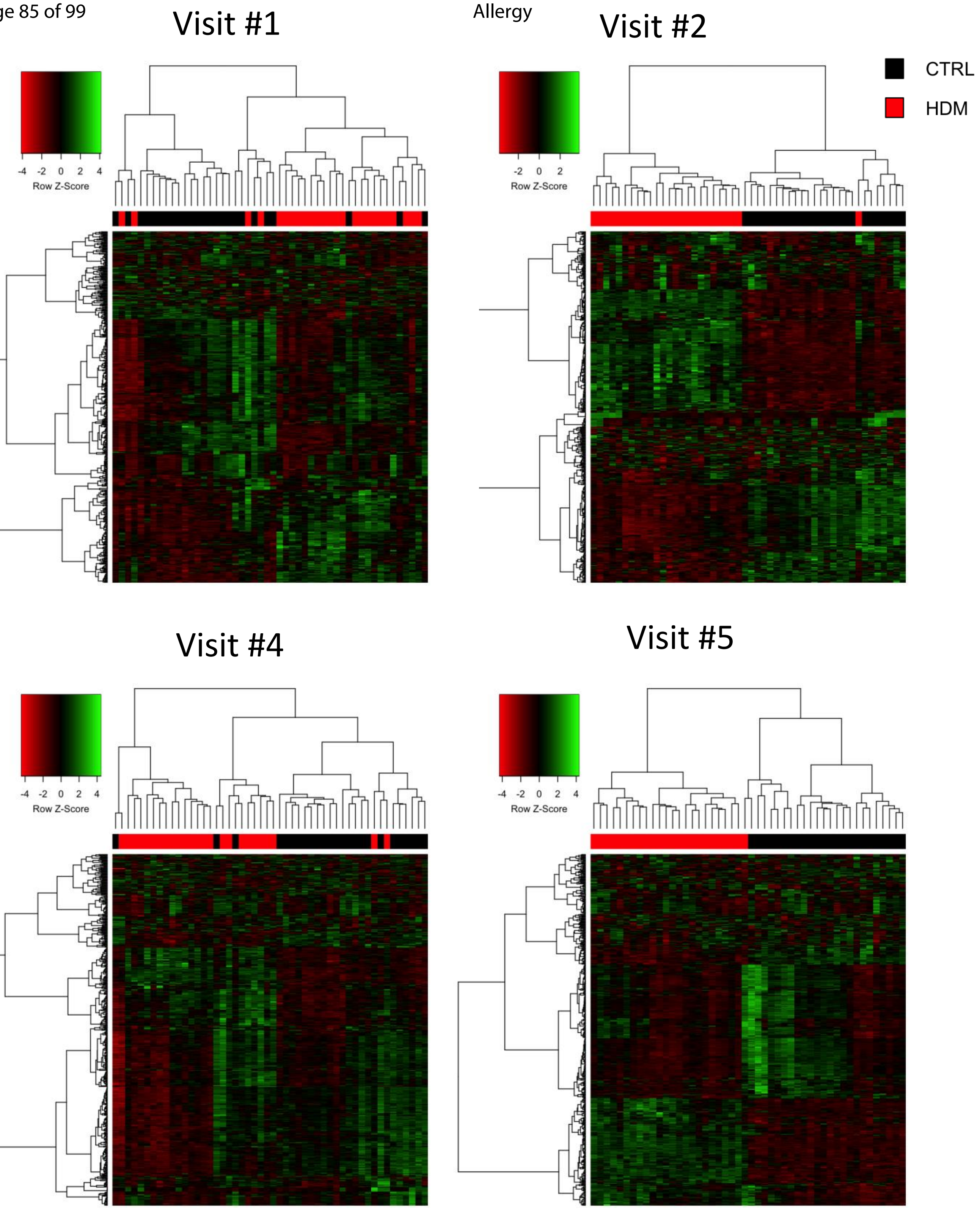


Supplementary Figure E6

Heatmap of *ComBat* adjusted data showing the 500 most variable genes for each individual patient (HDM/CTRL) at each time point (V1/2/4/5).

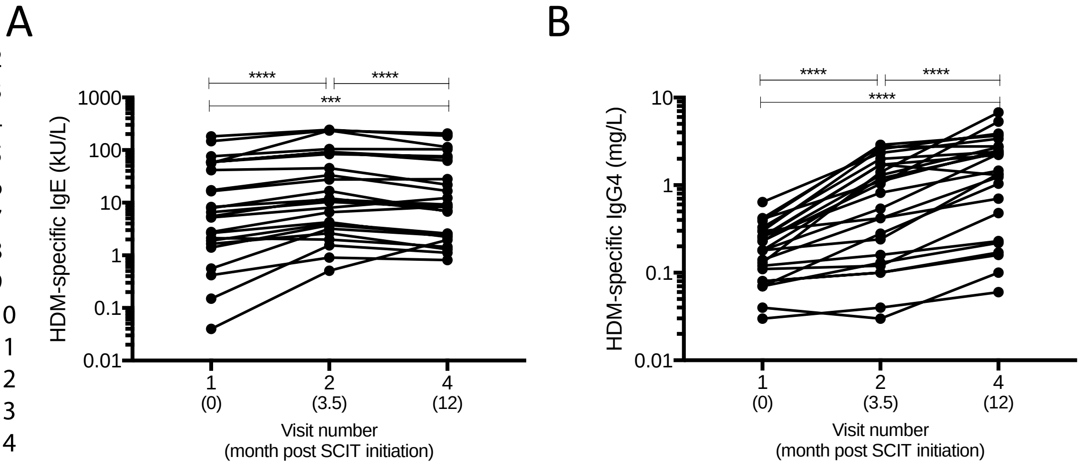


Supplementary Figure E7

### Supplementary Figure E8

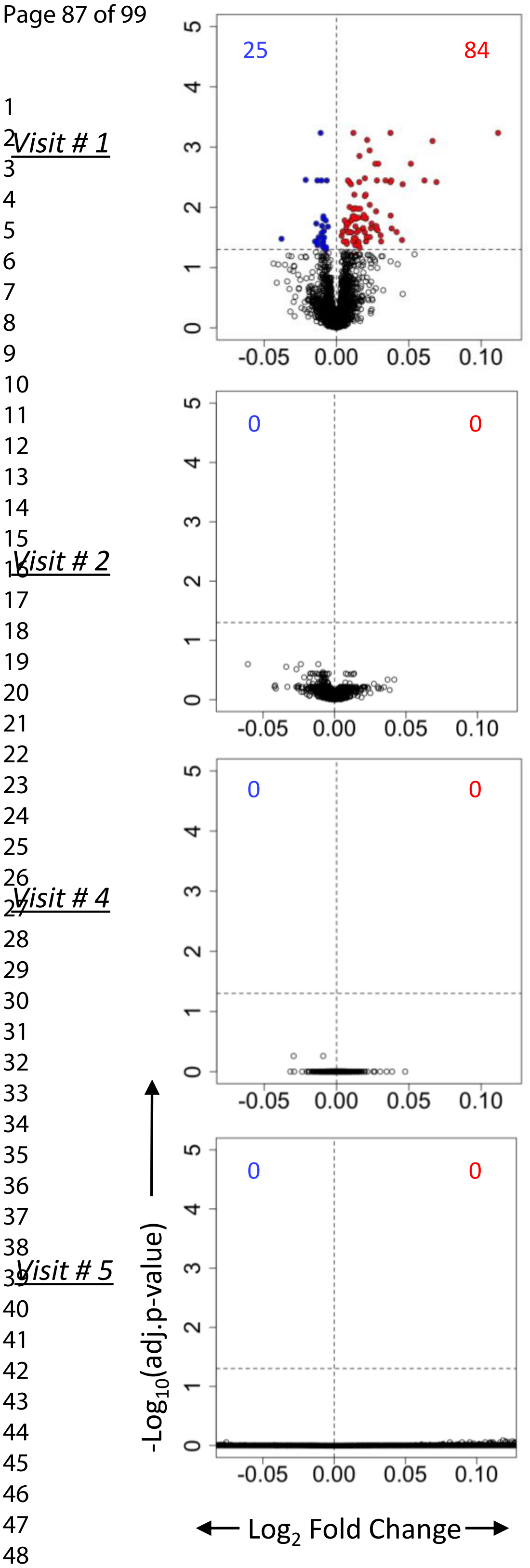


It was tested if response to stimulation is associated with respiratory symptoms at each visit employing *limma*. Volcano plots showing gene expression patterns associated cross-sectionally with attenuated post-treatment symptom scores at visits V1, V2, V4 and V5.

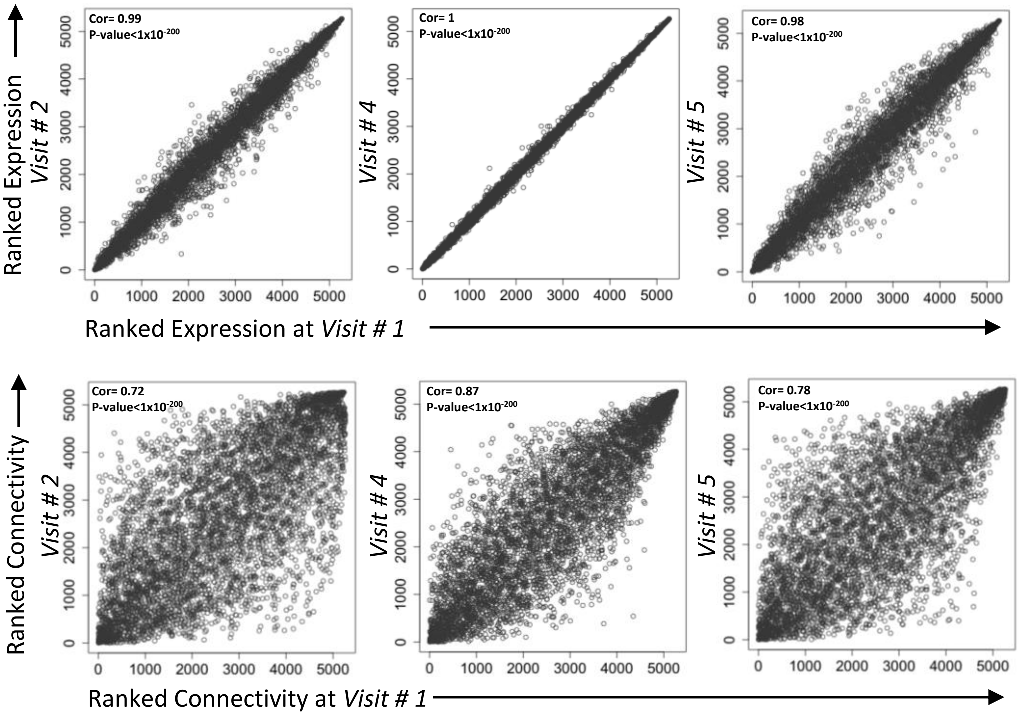


Supplementary Figure E9

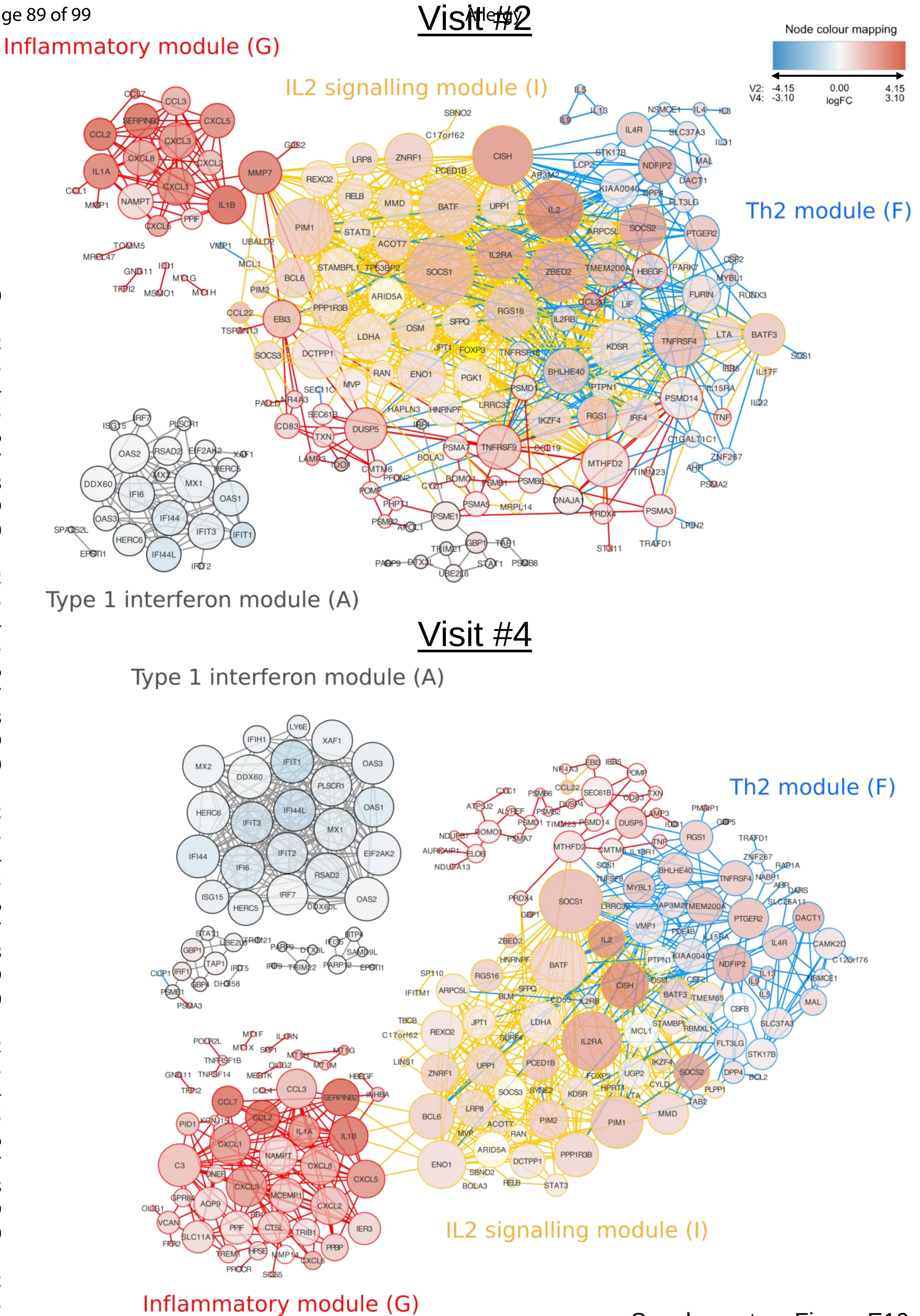


Supplementary Figure E10

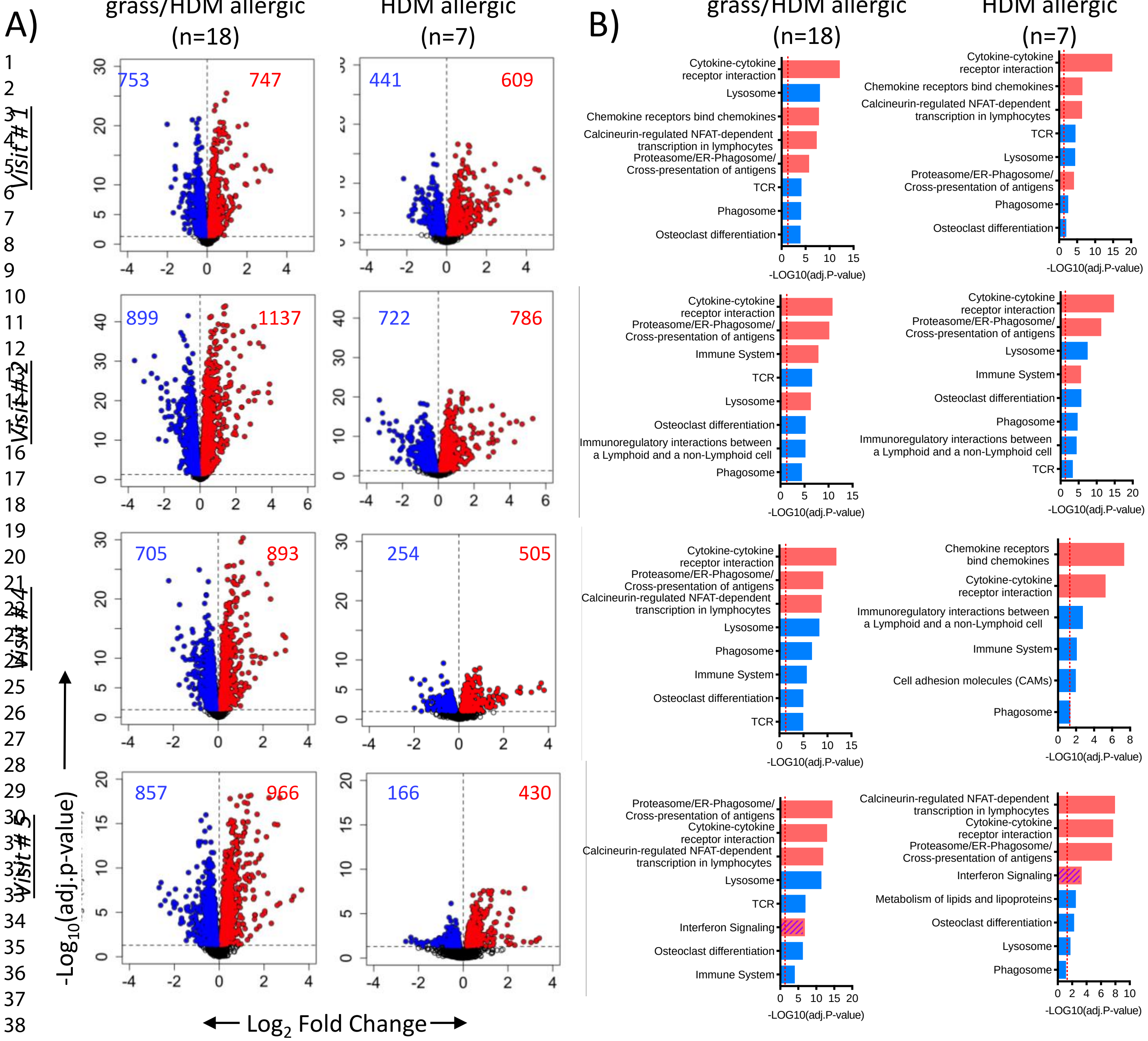


Supplementary Figure E11

A) Volcano plots showing differentially expressed genes in grass/HDM allergic (n=18) and HDM allergic (n=7) patients. B) Pathways analysis was carried out with InnateDB in grass/HDM allergic and HDM allergic patients.

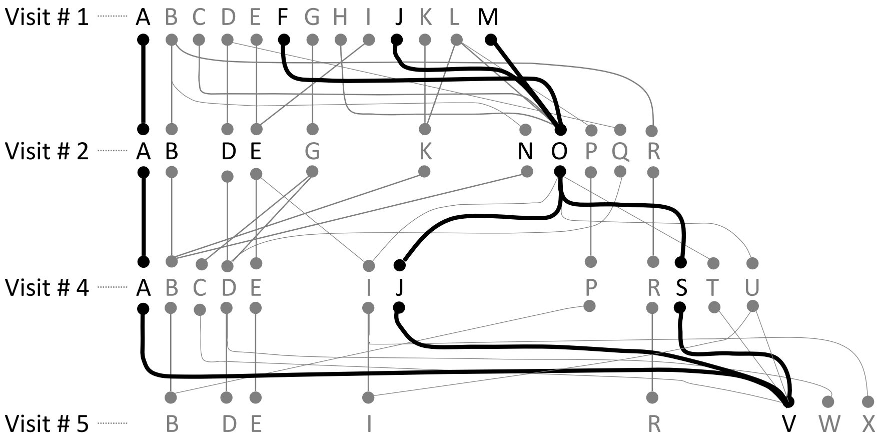


A = Type 1 interferon signalling
F = Th2
J = Inflammation/chemokine & cytokine receptor interactions/phagosome/proteasome
M = IL2 signalling
O = Th2, inflammation/proteasome/phagosome & cytokine receptor interaction, metabolism of proteins, IL2 signalling/Foxp3
S = Th2, proteasome/phagosome & cytokine receptor interaction, metabolism of proteins, IL2 signalling/Foxp3
V = Th2, inflammation/proteasome/phagosome & cytokine receptor interaction, metabolism of proteins, IL2 signalling/Foxp3, type 1 interferon

Supplementary Figure E12

Roadmap showing progressive re-wiring of the modules (A-M) over the course of SCIT in grass/HDM allergics.
